# Body reconstruction and size estimation of plesiosaurs: enlightenment on the ribcage restoration of extinct amniotes in 2D environments

**DOI:** 10.1101/2024.02.15.578844

**Authors:** Ruizhe Jackevan Zhao

**Affiliations:** Northwest University, Xi’an, China

## Abstract

Body size, especially body mass, is the key to understanding many biological properties. The scaling approaches and volumetric-density (VD) approaches are often employed to estimate the body masses of extinct amniotes. Precise skeletal reconstruction represents a pivotal step in all VD approaches, while the ribcage serves as one of the key determinants of thoracic shape and volume. Although being extensively investigated in physiological studies, the ribcage restoration remains poorly discussed during skeletal reconstruction. This study proposes one possible programme of skeletal reconstruction of extinct amniotes in 2D environments, focusing on the restoration of ribcage cross-sections. One recent VD approach, the cross-sectional method (CSM), was utilized to integrate the restored cross-sections into volume, therefore the workflow proposed here serves as a supplementary guideline of the application of the CSM in paleontology. Following this programme, a uniform set of reconstruction criteria was proposed for plesiosaurs, a clade of Mesozoic marine reptiles. Twenty-four plesiosaur models were created, then multiple regression models (Ordinary Least Squares, OLS; and Phylogenetically Generalized Least Squares, PGLS) based on them were employed to investigate the performance of various skeletal elements as size proxy. Despite the high disparity of their body plans, the trunk length and dimensions of dorsal vertebrae were found to be the most robost proxy for volume in plesiosaurs. The hybrid approach applied in this study, which incorporates VD estimates created under the same criteria as scaling samples, mitigates previous critiques focusing on inconsistent standards and inadequate taxonomic coverage. It allows fast and convenient body volume estimation for numerus individuals, even when only fragmentary fossil materials are available. The volumetric formulae for plesiosaurs can accommodate the size diversity of most taxa, except for some extremely giant pliosaurs, the largest of which might reach or exceed 20 metric tons in body mass. To demonstrate the utility of the formulae provided in this study, the body volumes of 113 plesiosaur taxa was estimated, and the branch-specific rates of size evolution computed from the data were mapped onto a plesiosaur phylogeny for visualization.

## Introduction

Body size has been long recognized as a pivotal indicator of many fundamental properties of an organism [1, 2, 3, 4]. The comprehension of size evolution in extinct animals not only illuminates the risks and opportunities they faced through deep time [5], but also establishes references for predicting the responses of extant species to environmental changes [6]. The growing requirement for a large amount of body size data is being echoed by the prevailing paradigm in plaeobiological research, which is reliant on machine learning tools and macroevolutionary models. For instance, phylogeny-based model selection enables the determination of the mode of size evolution within a specific clade [7, 8], while regression models between size and extrinsic factors provides clues for potential environmental drivers [9]. To collect sufficient amount of body size data, some previous studies employed one or multiple linear measurements of homologous structures as proxies [10, 11]. However, this method fails to account for potential errors arising from differences in body proportions, consequently hampering the comparison between taxa with distinct body plans. In contrast, body mass avoids the issue caused by proportional diversity, thereby serving as a more robust size metric.

Body mass estimation of extinct animals has always been a challenging task, primarily due to the fragmentary nature of the fossil record. To address this issue, the scaling approaches are often employed to estimate the dimensions of missing anatomical elements or the body masses directly [12, 13]. The workflow of the latter (namely extant scaling [ES] approaches [14]) is to establish regression equations between skeletal elements and body mass of extant species, and assumes that the same relationships hold for extinct clades [15, 16]. The advantage of the ES approaches is the computational efficiency, which allows rapid body mass estimation for numerous individuals [14]. But from a skeptical perspective, whether the predicted taxa (extinct) and the sampled ones (extant) follow the same rule remains agnostic. Such “worst” scenarios have been reported in the literature, exemplified by the 54 kg lower-bound mass estimate of extinct elephant *Anancus arvernensis*, despite its skeletal dimensions being comparable to adult African elephants *Loxodonta* [17]. Alongside with this risk, other cautionary issues imbedded in the scaling approaches have also been noted in previous studies (see [18] and citations therein).

Another category of method to estimate body mass is the volumetric-density (VD) approaches. The VD approaches involve various technical procedures to transform physical, 2D, or 3D models into volume estimates, which yield body mass metrics when multiplied by assigned body density [19, 20, 21]. Despite the recent advancement of 3D modelling techniques, its application in palaeobiology remains patchy in taxonomy due to the high investment demanded by accurate skeletal reconstruction and the difficulty in gaining access to fossil materials that scatter around the world. In contrast, the reduced time investment in 2D modelling facilitates widespread application of planar models and stimulates ongoing methodological advancement (e.g. [22, 23]). One of the latest 2D approaches was the cross-sectional method (CSM [24]). Compared with conventional methods requiring presumed cross-sectional geometries (e.g., ellipse [25, 20]), the CSM is capable of handling any shape and produces estimates with higher accuracy. The CSM defines the maximum height (or width) of a cross-section as the identidy segment, and the ratio of cross-sectional area to the square of identidy segment, *φ*, is introduced as a proxy for shape. After partitioning the model into multiple slices, the CSM assumes linear transitions for both shape and size of the cross-section at a small scale.

Skeletal reconstruction constitutes a pivotal step in all VD approaches [22, 26], while the ribcage serves as a critical determinant of thoracic cross-section. However, the ribcages of the fossil specimens are typically collapsed or incomplete, necessitating the estimation of the morphology *in vivo*. The ancestral state of rib design in amniotes was probably double-headed, consisting of a tuberculum and a capitulum that articulate with a diapophysis and parapophysis, respectively [27, 28]. Although such a plan is retained in most descendants, the dorsal ribs of some clades (e.g., squamates) are single-headed, being articulated to the diapophyses only [27]. It is also noteworthy that the ribcages of amniotes are mobile sturctures in life. While rib kinematics has been extensively investigated in physiological studies [29, 30], it remains poorly discussed in skeletal reconstruction. Most amniotes utilize costal aspiration to ventilate their lungs [31], whereby their ribs undergo periodical rotation synchronized with the respiratory rhythms. The rib kinematics can be decomposed into three compositions: bucket-handle rotation around a dorsoventral axis, pump-handle rotation around a mediolateral axis, and caliper rotation around a craniocaudal axis [28]. It has been argued that the morphology of costovertebral joints can reflect the rotational pathway of ribs to some extent [29], hence at least one static version of the rib orientation required by skeletal reconstruction can be inferred from the fossils.

Plesiosaurs were a clade of Mesozoic marine reptiles with volatile body plans, which were traditionally divided into two morphotypes: the long-necked, small-headed “plesiosauromorph” and the short-necked, large-headed “pliosauromorph” [32]. Later studies indicate that plesiosaurs exhibit a spectrum of body plans, and many species fall between the two endgroups [33]. Given the diversity in body proportions in plesiosaurs, total length is probably a poor indicator of body mass [34], while the performance of trunk length, which was typically utilized [35, 36], remains unverified. Although body sizes of plesiosaurs have attracted sustained scientific and public interest (e.g., the “25 m *Liopleurodon*” in *Walking with Dinosaurs*), body mass estimations for them remain scarce, predominantly emerging as byproducts of physiological studies [37, 38, 36, 39]. Moreover, the majority of plesiosaur models employed in published physiological studies were derived from fossil specimens with collapsed ribcages, museum mounts with more than a century of history, or outdated reconstructions in historical literature. The only study focusing on size estimation to date was [34], in which the body lengths of some thalassophonean pliosaurs were calculated. The body volumes, however, were based on a commercial model, overlooking the interspecific disparity in body proportions. While multiple plesiosaur skeletal reconstructions were presented in one popular science publication [40], the corresponding body mass estimation procedure was inadequately described, probably due to considerations for readership.

Although body mass estimations based on rigorous skeletal reconstructions are scarce in plesiosaurs, the cumulative contribution of previous studies has established a feasible fundation for this attempt. Some plesiosaur possessed the highest number of cervical vertebrae among all vertebrates [41, 42], which attracted particular research attention. Comprehensive vertebral measurements of multiple plesiosaur specimens are available in the literature (e.g. [43, 44]), and in turn enable reliable body length estimations. The paradigm of body length estimation proposed for pliosaurs in [34] is also applicable to all plesiosaurs (and many other extinct amniotes as well). The diversity of rib orientations in different plesiosaur taxa and their implications for body shape variations have been noticed in previous studies [45]. Furthermore, the preservation of soft tissue impressions in multiple plesiosaur fossils provides extra insights to their body shapes *in vivo* [46, 47, 48, 49].

Some studies adopted a hybrid approach to estimate body masses of extinct amniotes, by developing regression formulae based either on VD estimates or composite datasets incorporating both extinct and extant taxa (see [14] for a comprehensive review of the advantages and critiques mentioned below). The hybrid approaches can mitigate the limitations of both VD and ES approaches: the substantial time investment in modelling and the unverified applicability of the formula, respectively. One major critique of the hybrid paradigm concerns the inconsistent reconstruction standards underlying the selected VD estimates, and taxonomic biases during sampling. Ideally, if a unifrom set of criteria is proposed for a specific clade of extinct amniotes, and implemented in multiple representative taxa, the penalty of this shortcoming would be reduced. In this paper, I present one possible protocol for 2D reconstruction of plesiosaurs, and created 24 models under these criteria. The performance of various skeletal structures as size proxy was evaluated using regression models, and the body masses of 113 plesiosaur taxa were estimated to demonstrate how this paradigm facilitates downstream analyses. The 2D reconstruction methodology employed in this study, particularly the ribcage restoration protocol, can be extended to many clades of extinct amniotes, therefore works as a supplementary guideline of the application of the CSM in paleontology.

## Results

### 2D skeletal reconstruction of extinct amniotes

The modelling workflow employed in this study is presented here as a research result, establishing a programme of precise skeletal reconstructions of extinct amniotes in 2D environments (see Materials and Methods for its detailed application in plesiosaurs). However, it is not considered a dogmatic solution, but an invitation for future advances in reconstruction techniques. As noted in the Introduction, the scaling approaches are typically employed to estimate the sizes of unpreserved skeletal elements, represented by regression equations that reveal their relationships within a specific clade, and comparisons with phylogenetically proximate taxa posessing more complete fossil remains (assuming uniform body proportions and absence of allometry; Fig. 1). The main body axis is first created, incorporating all anatomical elements contributing to the total body length, including the skull, vertebrae (and intercentra if present), and intervertebral cartilage. Previous studies have emphasized the critical role of vertebral reference sets [34]: when the spine is incompletely preserved in a specimen, its body length along the vertebral column (excluding intervertebral cartilage) can be estimated by comparing vertebral dimensions with those of phylogenetically or functionally proximate individuals possessing complete vertebral sequences. Intervertebral cartilage is also an important component of body length, the dimensions of which should be inferred from articulated fossils. It has been revealed that intervertebral distance increase congruently with vertebral dimensions, rendering proportional values a more reliable quantitative standard than sheer sizes [50]. The restoration of spinal curvature can be regarded as an integral step in ribcage reconstruction, both of which are elaborated below. In the 2D reconstructions followed by the CSM, all limbs are created separately and treated as discrete objects to facilitate volumetric calculations. Soft tissue reconstruction constitutes the final modelling step, which is frequently constrained by the fossil materials, with only exceptionally preserved specimens providing relavant information. It is noteworthy that even in nearly complete fossils, the collapse of ribcage may introduce uncertainty while interpreting the thickness of soft tissues (see Discussion).

**Figure 1:**
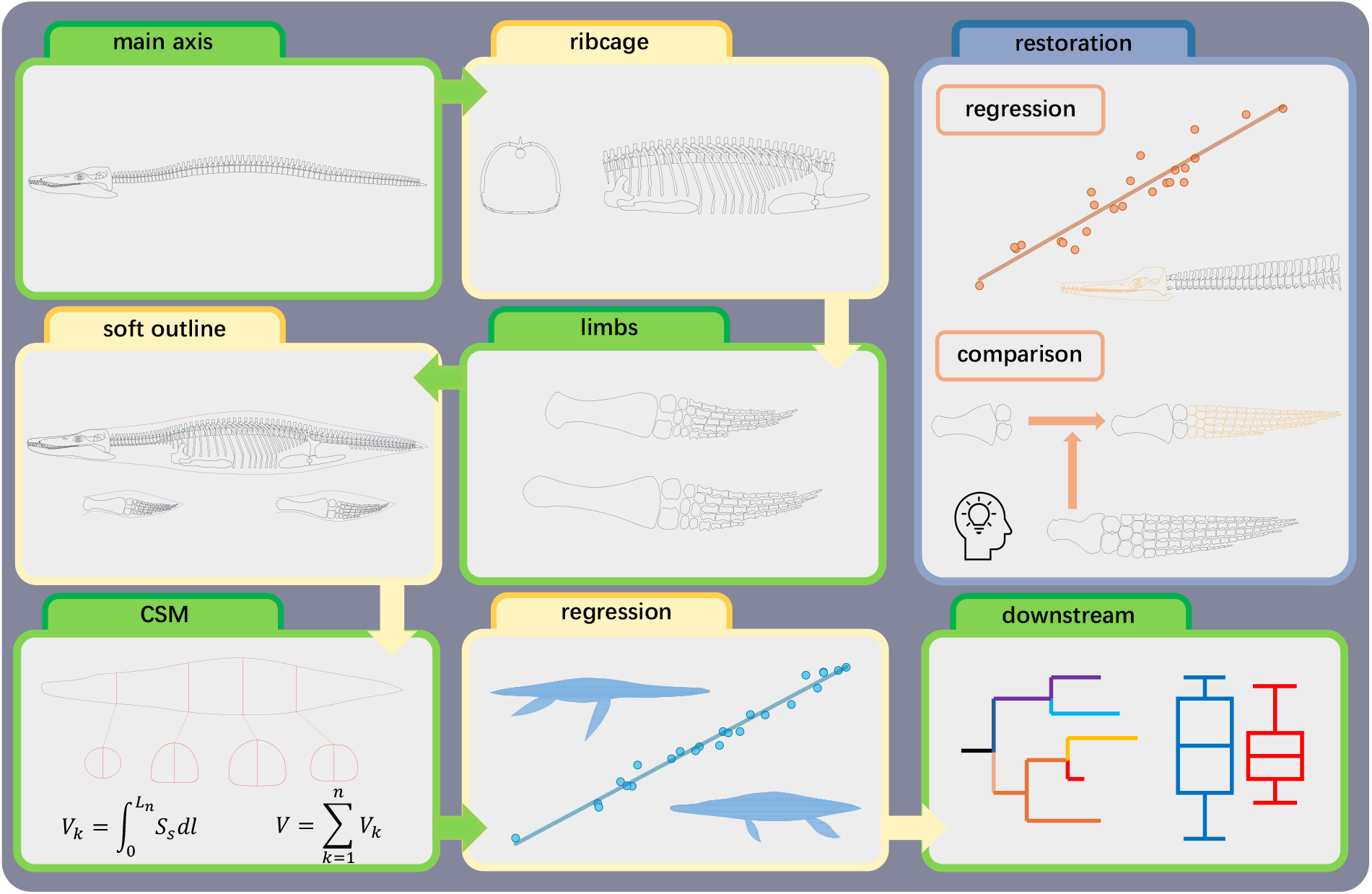
Workflow of the reconstruction programme proposed in this study. All components are schematic representations: skeletal elements and silhouettes are not to scale, and regression lines and downstream models don’t necessarily correspond to those used in this study.

Spinal curvature serves as a pivotal indicator for body shape in many clades of amniotes, thus restoration is demanded for fossil specimens lacking articulated vertebral columns. The mathematical model based on wedging-angles, initially developed for primates, has been successfully applied to a few plesiosaur taxa before [51]. Nevertheless, this approach mandates complete and disarticulated vertebral series free from taphonomic distortion, coupled with plural forms of measurements. Such requirements are seldomly fulfilled throughout the fossil record, thus a compensatory approach is proposed here. The vertebral columns of most amniotes exhibit pronounced regionization (e.g., cervicals, dorsals [52, 53, 27]). If an anatomical correspondence between the vertebral regions and the girdle elements is established, the spinal curvature can be inferred given the spinal lengths of different regions and the inter-girdle distance.

To infer potentail rib orientation from the dorsal vertebrae, three angles are introduced here (Fig. 2A): angle *θ*_1_ ∈ [− 90°, 90°] quantifies the deviation of pump-handle rotation from the vertical plane, inferred from the diapophysis or the axis formed by the diapophysis and the parapophysis; angle *θ*_2_ ∈ [0°, 90°] quantifies the deviation of bucket-handle rotation from the vertical plane, inferred from the diapophysis; angle *θ*_3_ ∈ [−90°, 90°] quantifes the deviation of vertebral centrum from the horizontal plane. It is noteworthy that angles *θ*_1_ and *θ*_2_ may not remain the same throughout the dorsal region. For instance, the transverse processes of plesiosaurs indicate a gradual change in rib orientation through the vertebral column (Fig. 2B), which was described in multiple specimens [54, 45] but neglected in some published skeletal reconstructions (e.g., [40]).

**Figure 2:**
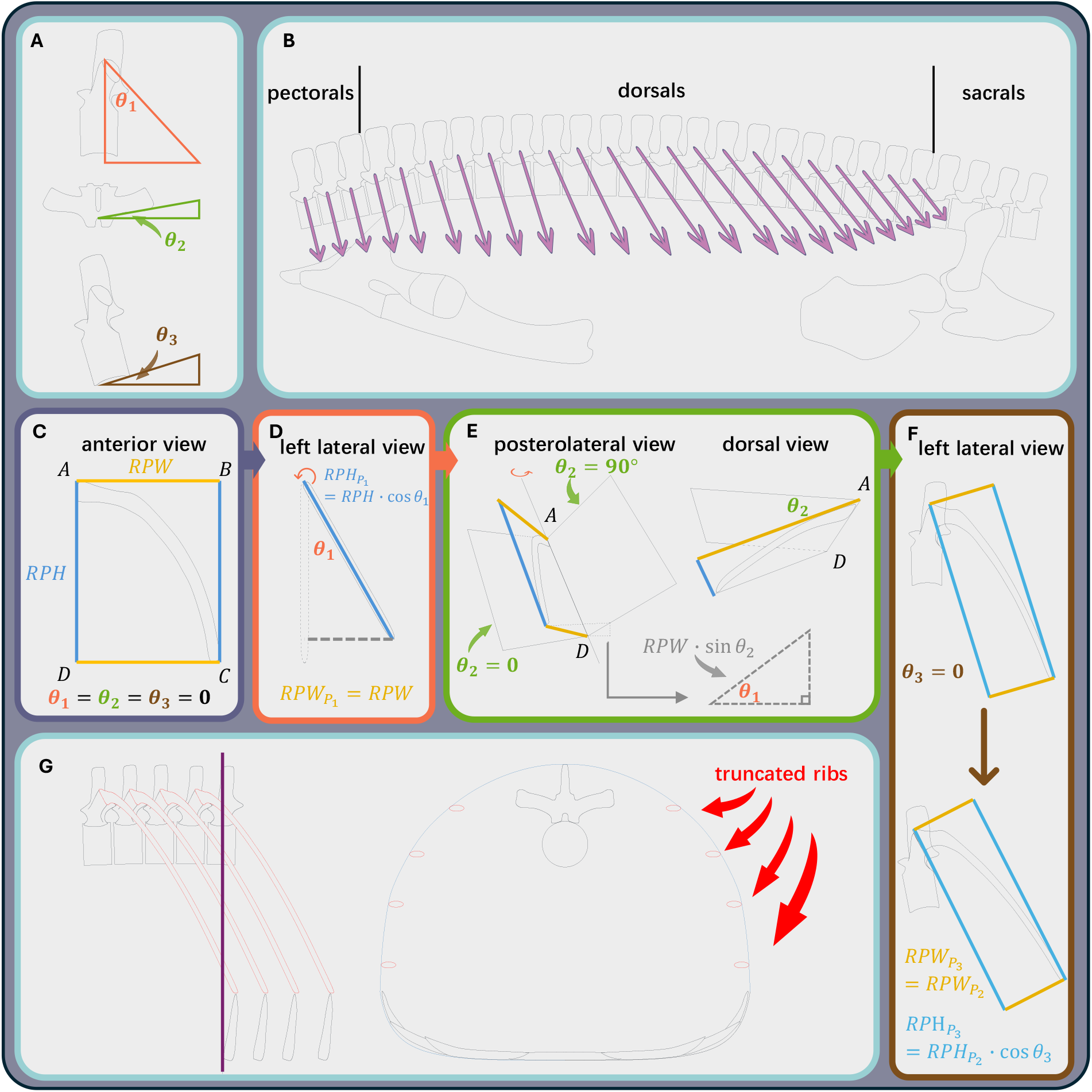
Protocol of the rib rotation (*θ*_1_ → *θ*_2_ → *θ*_3_) and ribcage restoration. (A) The method to infer angles *θ*_1_, *θ*_2_, and *θ*_3_ from the vertebrae. (B) A schematic ribcage of plesiosaur *Cryptoclidus eurymerus* showing the gradual change in angle *θ*_1_ from lateral view, angles inferred from the restored vertebral column provided in [51]. (C) The vertically orientated rib plane (*θ*_1_ = *θ*_2_ = *θ*_3_ = 0) showing the rib plane height (RPH) and rib plane width (RPW). (D) Lateral view of the rib plane that rotates around the AB axis by *θ*_1_ degrees. (E) Posterolateral and dorsal views of the rib plane that rotates around the AD axis by *θ*_2_ degrees. (F) Lateral view of the costovertebral system that rotates around the mediolateral axis by *θ*_3_ degrees. (G) Schematic images showing the method to determine the dorsal outline of the ribcage from truncated ribs.

The dorsal rib is first simplified as a 2D structure, and assumed to lie within a vertical plane (Fig. 2C). The minimum rectangle containing the rib is drawn and reveals its height and width (termed here as the rib plane height [RPH] and the rib plane width [RPW]). The four vertices of the rectangle is labeled A, B, C, and D sequentially. Angle *θ*_1_ is considered first, and the rib plane is rotated around the AB axis by *θ*_1_ degrees (Fig. 2D). Then RPH projected to the vertical plane in this moment 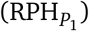 equals RPH *·* cos *θ*_1_, while the projected RPW 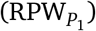 equals RPW. Then the rib plane is flipped around the AD axis by *θ*_2_ degrees (Fig. 2E), and the projected height 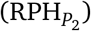 and width 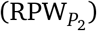 are given by:

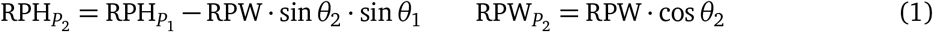

After the determination of rib orientation inferred from the vertebral morphology, the costovertebral system is rotated around the mediolateral axis by *θ*_3_ degrees (Fig. 2F). Then the projected height and width are 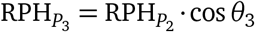 and 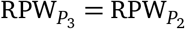, respectively. Subsequent to the reconstruction of the ribcage in lateral view, a specific vertical plane is drawn to truncate multiple dorsal ribs (Fig. 2G). The truncated width of each rib plane (RPW_*i*_, *i* = 1, 2, · · ·) is then determined, the projected width (RPW_*iP*_) equals RPW_*i*_ ·cos *θ*_2_. The dorsal outline of the ribcage formed by the ribs is then created by connecting the projected truncation points with smooth curves. It can be complemented into a cross-sectional profile if the ventral outline (e.g., of the gastralia if present) is assumed.

It is noteworthy that 3D rotation is not commutative. In other words, prioritizing either angle *θ*_1_ or angle *θ*_2_ would result in different final positions of the ribs, thereby leading to uncertainty in ribcage reconstruction. To quantify its impact, each plesiosaur model produced in this study was subject to a sensitivity test (i.e., angle *θ*_2_ being rotated first, followed by angle *θ*_1_, Fig. S1; the mathematical model is provided in the Supplementary Text). However, it is proposed here that prioritizing angle *θ*_1_ in amniote reconstruction has advantages in both theory and practice (see Discussion).

### Regression equations

A series of regression equations was established to estimate the skull length (SKL), maximum rib arc length (max RAL), trunk length, and tail length of plesiosaurs, respectively (all linear measurements are in millimeters, see Fig. S2 for the measuring standards). The samples of all regression models are phylogenetically widespread, containing species from every major clade of plesiosaurs (i.e., Rhomaleosauridae, Pliosauridae, Microcleididae, Cryptoclididae, Leptocleididae, Polycotylidae, Elasmosauridae), so that these models are probably applicable to the whole Plesiosauria. In case of type I errors caused by shared evolutionary history, Phylogenetic Generalized Least Squares (PGLS) was used to reveal the size correlation among skeletal elements. The PGLS models are performed on the pruned maximum clade credibility tree (MCCT) summarized from the post burn-in Bayesian tree samples (see Materials and Methods), which is consistent with published trees in general topology ([55, 56]; Fig. S3). Ordinary Least Squares (OLS) were also applied on the same datasets, and their performance was compared with that of corresponding PGLS models by sample-size corrected Akaike

Information Criterion (AICc) values and mean per cent prediction errors (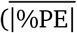 [57], see Materials and Methods for definition). While the *R*^2^ value quantifies the explanatory power of the independant variable to the dependant variable, the 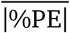 value evaluates the prediction performance of the regression model. For the trunk-rib (N = 24) and trunk-tail (N = 19) datasets, the PGLS models reveal significant correlation between selected variables (*P* < 0.001), and the OLS models yield lower AICc values. The 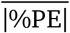 values, on the other hand, are slightly lower in both PGLS models than the corresponding OLS:

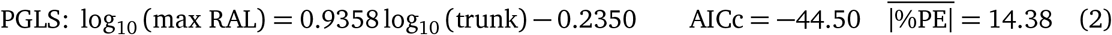

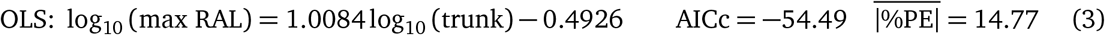

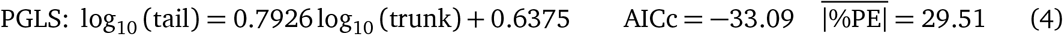

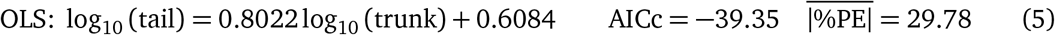

In general, the PGLS models fitted to the two datasets demonstrate worse fitness than the OLS judging from the AICc values (as in some previous studies [7, 14]) and negligible advantage in prediction (<1%). Given that future revisions of the phylogeny may also affect the PGLS results, the OLS models (*R*^2^ = 0.8973 in the trunk-rib regression; and 0.7808 in the trunk-tail regrssion) were employed in this study.

An extra nonlinear regression based on a log-logistic (LL) function was fitted to the skull-neck dataset, in addition to PGLS and OLS (see Discussion for rationale). The PGLS model also reveals a significant correlation between the skull-neck ratio and the cervical number (CN; *P* < 0.001). Among the three models, the nonlinear model significantly outperformes both the PGLS and OLS models, evidenced by the lowest AICc value and 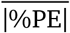 value. It was thus used in subsequent reconstructions due to its performance in both fitness and prediction.

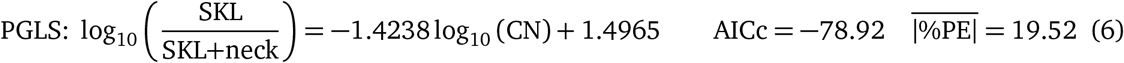

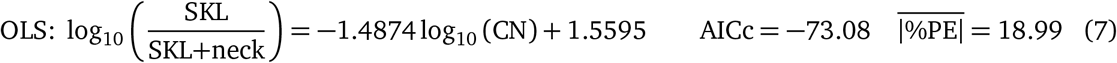

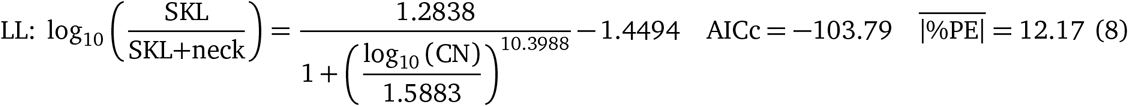

### Plesiosaur models and volume equations

Twenty-four plesiosaur models were created following the reconstrution criteria proposed in this paper (see Materials and Methods). The body sizes of some thalassophonean pliosaurs and elasmosaurids dwarf those of other plesiosaur clades, as shown in figure 3. The estimated body lengths along the vertebral column and body volumes are prsented in Table 1. Assigning a uniform body density is equivalent to multiply all volumetric estimates by a constant coefficient while the relative size relationships remain unaltered. Given the daily variation in body density of aquatic tetrapods [58] and the habitat diversity of plesiosaurs (pelagic vs fresh water [59]), volumetric estimates rather than body masses were incorporated into the regression analyses, thereby maintaining adjustable density for future studies.

**Table 1:** Estimated body lengths and volumes of the 24 plesiosaur models. Osteologically immature individuals are marked with a *. See the Supplementary Text for institutional abbreviations.

| Taxon | Catalogue Number | Clade | Length (m) | Volume (m<sup>3</sup>) |
| --- | --- | --- | --- | --- |
| *Albertonectes vanderveldei* | TMP 2007.011.0001 | Elasmosauridae | 12.122 | 3.0098 |
| *“Aristonectes” quiriquirensis* | SGO.PV.957 | Elasmosauridae | 10.000 | 4.0562 |
| *Hydrotherosaurus alexandrae* | UCMP 33912 | Elasmosauridae | 8.794 | 1.8277 |
| *Styxosaurus* sp. | SDSM 451 | Elasmosauridae | 11.254 | 2.9987 |
| *Thalassomedon hamingtoni* | DMNH 1588 | Elasmosauridae | 11.953 | 6.6125 |
| *Vegasaurus molyi* | MLP 93-I-5-1 | Elasmosauridae | 6.168 | 0.8172 |
| *Abyssosaurus nataliae* | MChEIO PM/1 | Cryptoclididae | 6.739 | 1.2057 |
| *Cryptoclidus eurymerus* | NHMUK PV R2860 | Cryptoclididae | 3.753 | 0.3613 |
| *Dolichorhynchops osborni* | FHSM VP404 | Polycotylidae | 2.958 | 0.4137 |
| *Martinectes bonneri* | KUVP 40002 | Polycotylidae | 4.947 | 1.2330 |
| *Mauriciosaurus fernandezi* * | INAH CPC RFG 2544 PF1. | Polycotylidae | 2.114 | 0.0776 |
| *Polycotylus latipinnis* | YPM 1125 | Polycotylidae | 5.073 | 1.0292 |
| *Kronosaurus/Eiectus* | MCZ 1285 | Pliosauridae | 10.364 | 11.0968 |
| *Liopleurodon ferox* | GPIT-RE-3184 | Pliosauridae | 5.825 | 1.7482 |
| *“Monquirasaurus” boyacensis* | MJACM 1 | Pliosauridae | 9.080 | 9.6561 |
| *Peloneustes philarchus* | GPIT-RE-3182 | Pliosauridae | 4.337 | 1.0075 |
| *Pliosaurus* cf. *kevani* | CAMSM J.35990 | Pliosauridae | 9.714 | 10.4712 |
| *Pliosaurus funkei* | PMO 214.135 | Pliosauridae | 9.714 | 10.5937 |
| *Sachicasaurus vitae* | MP111209-1 | Pliosauridae | 10.396 | 12.2288 |
| *Stenorhynchosaurus munozi* * | VL17052004-1 | Pliosauridae | 5.951 | 1.8211 |
| *Macroplata tenuiceps* | NHMUK PV R5488 | Rhomaleosauridae | 4.627 | 0.6703 |
| *Meyerasaurus victor* | SMNS 12478 | Rhomaleosauridae | 3.391 | 0.3639 |
| *Microcleidus tournemirensis* | MMM J. T. 86-100 | Microcleididae | 4.210 | 0.2173 |
| *Seeleyosaurus guilelmiimperatoris* | SMNS 12039 | Microcleididae | 3.658 | 0.1940 |

**Figure 3:**
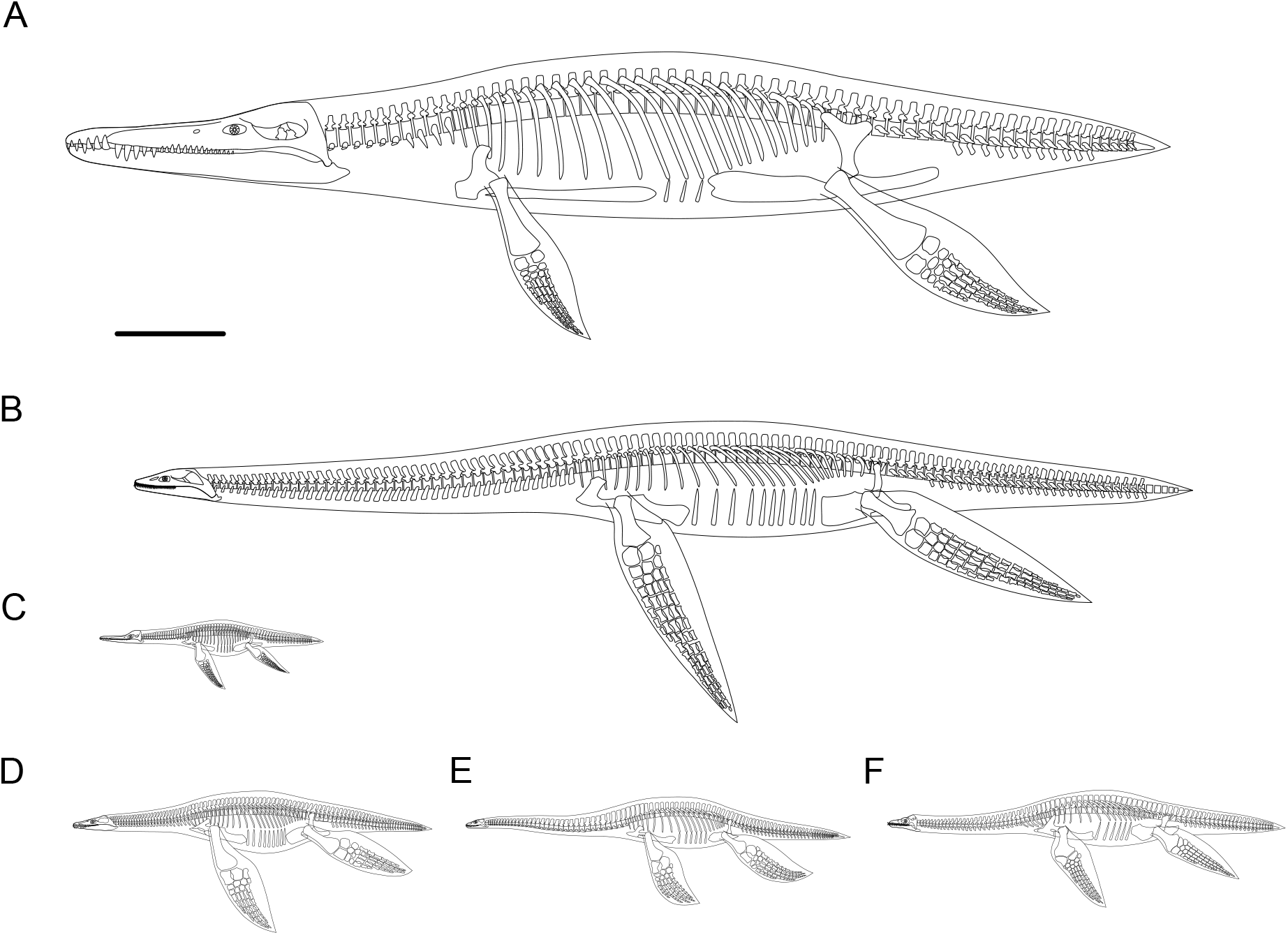
Representative plesiosaur models from different clades. (A) *Kronosaurus/Eiectus*. (B) *“Aristonectes” quiriquinensis*. (C) *Mauriciosaurus fernandezi*. (D) *Meyerasaurus victor*. (E) *Seeleyosaurus guilelmiimperatoris*. (F) *Cryptoclidus eurymerus*. All models were constructed under the *θ*_1_ → *θ*_2_ rib rotational sequence. All limbs are vertically orientated for display. Scale bar equals 1 m.

Table 2 shows the performance of 8 skeletal structures as proxy for body volume tested using OLS, and the PGLS results based on the same datasets are available in Table S1. Given the diversity in body proportions among plesiosaurs, the total body length and skull length were not subject to regression tests. All tested variables demonstrated significant correlations with estimated volumes (P<0.001 in PGLS models). Among all proxies, the trunk length and the dimensions of dorsal vertebrae (DDV; defined as mean length ° mean width ° mean height of the dorsal centra) yield the best performance, both showing high *R*^2^ values and low 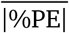 values. Similar to the regression models for rib and tail, the PGLS versions of trunk-volume and DDV-volume formulae don’t show significantly lower AICc values (|*Δ*AICc| *>* 2) than the OLS models, hence the later was recommended for predictive applications. Different from the confidence intervals, the 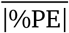 value provides a symmetrical error range when the estimated volumes are back-transformed to its antilog scale [14]:

**Table 2:** Parameters of the regression models based on ordinary least squares (OLS). All countinous measurements are in meters (m), and all proxies and the volumes were log_10_-transformed before analysis. Abbreviations: N, sample size; %PE, mean per cent prediction error; SKL, skull length; CN, cervical number; DDV, dimensions of dorsal vertebrae (mean length ° mean width ° mean height of the dorsal centra). See Fig. S2 for measuring criteria.

| Proxy | Slope | Intercept | N | P-value | $R^2$ | $\vert \%PE\vert $ |
| --- | --- | --- | --- | --- | --- | --- |
| SKL×CN | 2.9233 | -3.5193 | 16 | <0.001 | 0.8802 | 56.15 |
| trunk | 2.8223 | -0.5317 | 24 | <0.001 | 0.9831 | 16.85 |
| DDV | 1.0224 | 3.5117 | 14 | <0.001 | 0.9770 | 19.08 |
| humerus length | 2.7906 | 1.1966 | 23 | <0.001 | 0.7827 | 59.94 |
| humerus chord | 3.3229 | 2.3354 | 23 | <0.001 | 0.7725 | 67.26 |
| femur length | 2.4673 | 1.0266 | 21 | <0.001 | 0.7604 | 74.45 |
| femur chord | 2.9865 | 2.1182 | 21 | <0.001 | 0.8708 | 47.92 |
| coracoid length | 2.5371 | 0.7703 | 16 | <0.001 | 0.8909 | 51.87 |
| coracoid width | 2.7213 | 1.982 | 22 | <0.001 | 0.8917 | 42.08 |
| pubis length | 2.2420 | 1.0138 | 17 | <0.001 | 0.8366 | 56.48 |
| pubis width | 2.9977 | 1.4618 | 19 | <0.001 | 0.9154 | 43.94 |
| ischium length | 2.0481 | 0.9642 | 19 | <0.001 | 0.7999 | 65.52 |
| ischium width | 3.0514 | 1.8605 | 19 | <0.001 | 0.9372 | 40.88 |

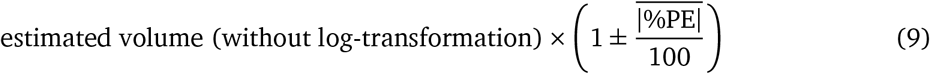

To evaluate the impact of rib rotational sequence on volumetric estimation, an alternative *θ*_2_ →*θ*_1_ version was created for each plesiosaur model to conduct a sensitivity test. As shown in Table S2, the rotational sequence caused limited influence on the total body volume, with the differences of all models lower than 3.5% and the average difference being 1.27%.

### Rates of body size evolution in plesiosaurs

To demonstrate how the methodological framework established in this study facilitates volumetric estimation, the body volumes of 113 plesiosaur taxa out of 146 in the current phylogeny were estimated, based on skeletal model, trunk length, DDV, ischium width, femur chord, or SKL×CN. The dataset is more than 2 times larger than the one containing 48 plesiosaur taxa used in a recent study [36]. Figure 4A shows the branch-specific rates of volumetric evolution mapped onto the pruned MCCT. Given the unsatisfactory performance of some size proxies, the dataset was reduced to a subset that solely contains the volumes of osteologically mature individuals estimated from models, trunk length, or DDV. Inferred evolutionary rates in body volume mapped onto the MCCT is presented in Fig. 4C. The estimated evolutionary rates were log_10_-transformed before mapping since extremely high values in specific branches may conceal the overall pattern (i.e., most branches will exibit cold colors). The comparison of a heterogeneous model against a homogeneous one was replicated on another 100 trees randomly selected from the post burn-in Bayesian samples (see Materials and Methods). In alignment with the result discovered in [36], the heterogeneous model outperformes the homogeneous model, evidenced by the log Bayes Factors (BF; positive evidence [log BF*>* 2] in 92 trees; strong evidence [log BF*>* 5] in 71 trees; very strong evidence [log BF*>* 10] in 25 trees).

**Figure 4:**
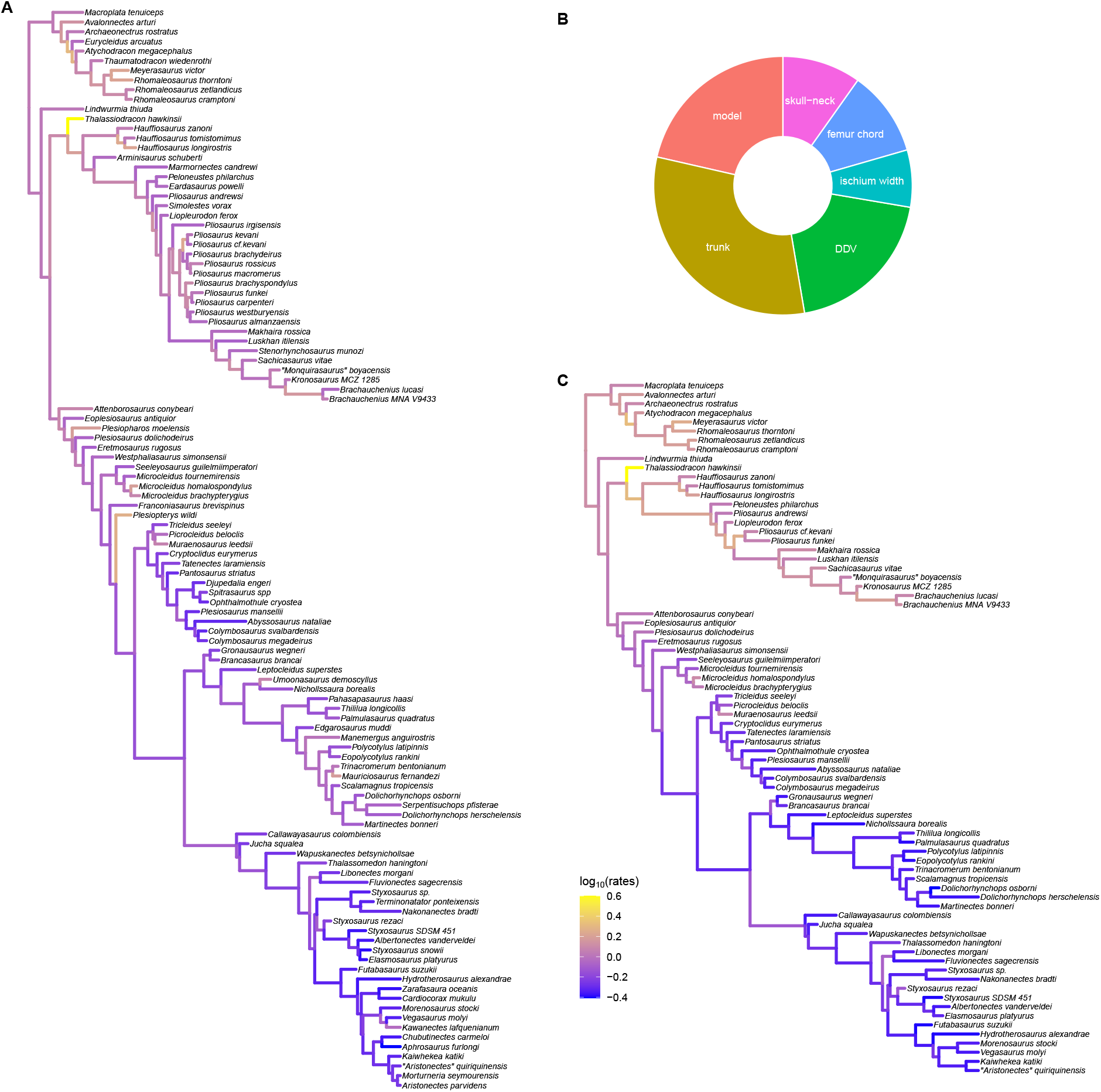
Analytic results of plesiosaur body size evolution. (A) Pruned MCCT with the size evolutionary rates of 113 plesiosaur taxa mapped onto the branches, including osteologically immature individuals. (B) Pie chart showing the data sources of body mass estimates presented in (A), and their proportions within the dataset. See the main text and Fig. S2 for the definitions of size proxies. (C) Pruned MCCT with the size evolutionary rates mapped onto the branches. Only osteologically mature individuals estimated from the model, trunk length, or DDV are included.

## Discussion

### Non-linearity after log-transformation in skull-neck relationship

Linear regression models based on OLS can effectively capture the covaration of two sets of linear measurements if they follow an isometric relationship. Biological systems, however, frequently exhibit power-law scaling patterns [60]. Geometrically, doubling the length of an object will increase its volume eightfold. Logarithmic transformation of such datasets can convert the power-law relationships to the ones that can be fitted with linear functions. However, the typical assumption that log-transformation completely linearises the relationship is not always satisfied [18]. Some previous studies reported the potential systematic biases in prediction arising from this assumption, revealing that nonlinearity after log-transformation is not uncommon in biological scaling [61, 62]. However, neither well-defined criteria for determining when to apply nonlinear regression nor the models incorporating phylogeny have been developed. Therefore, the application of nonlinear models relies entirely on empirical evaluation in the current stage.

The distribution of sample points of the log-transformed skull-neck data is shown in Fig. S4. When a linear model is fitted, the data points at both ends of the horizontal axis tend to fall below the regression line, while those in the middle lie above the line, stimulating the author’s intuitive doubt about the validity of linear fitting. The distribution of scatter points geometrically mimics the graph of a logistic curve, hence the dataset was subject to a nonlinear regression using a log-logistic function. The AICc and 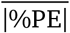 values indeed indicate that this model yields better performance than the OLS or PGLS regression, thereby it was employed in estimating the SKLs of plesiosaur models in this study. The scatter plots of all other regression models were also manually examined, and no nonlinear model after log-transformation was considered feasible for them.

### Uncertainty in soft tissue reconstruction of plesiosaurs

The quantitative restoration of soft tissues, encompassing muscles, fat and skin, constitutes a pivotal step in all VD approaches [63, 21]. It helps determine both volume and body shape of an extinct amniote, thereby creating a substantial impact on downstream physiological studies. The fossil record should be regarded as a prioritised reference when soft tissue impressions are preserved. However, such luxurious cases are extremely scarce, and there exists potential discrepancy in the interpretation of soft imprints as evidenced by fossil specimens, as shown below. Information from extant analogues is thus frequently utilised to address these knowledge gaps [21, 64]. Soft tissue impressions are preserved in some plesiosaur fossils, with the earliest reports tracing back to the 19th century [65]. The majority of plesiosaur fossils with soft tissue traces are microcleidids from German deposites, with soft tissue remains predominantly surrounding the flippers or the tail [66, 49, 48]. *Mauriciosaurus fernandezi*, on the other hand, is known from an almost complete specimen with extensive soft tissue in the trunk region, illuminating the body outline of plesiosaurs *in vivo* [46]. However, the existence of these fossils does not solve the problem of plesiosaur reconstruction once and for all. There are currently no plesiosaur fossils that reveal the soft contours of the leading edges of the flippers or the propodial regions, and the shape of the flipper *in vivo* was probably not a direct extension of the convex hull formed by the limb bones, according to modern aquatic tetrapods [67]. Fossil evidence for the amount of soft tissue on the dorsal and ventral sides of the ribcage in plesiosaurs is also absent, necessitating the introduction of modern analogues to complete this puzzle. The following review is presented on the subject of soft tissue restoration in plesiosaurs, involving established muscular restoration, physiological knowledge, and associated information from extant aquatic tetrapods, in order to provide theoretical support for the reconstruction criteria proposed in this study. However, it should be alerted that when extrapolating the body plans of extinct species according to extant ones, the underlying assumption involves convergent evolution of particular biological structures driven by natural selection. Therefore the inference may carry the risk of over-reliance on adaptationist explanations [68].

The soft contour at the skull region of plesiosaurs has not been quantitatively discussed in previous literature. It has been argued by several researchers that a boss-like structure on the ventral symphyses of certain elasmosaurid mandibles may be the attaching site of the geniohyoid muscle [69, 70, 71], offering some limited clues for the cranial profile. For instance, its morphology in *“Aristonectes” quiriquinensis* indicates a loose mouth floor [72], while the general outline remains unknown. Among extant amniotes, aquatic mammals generally possess well developed craniofacial soft tissues, which function as echolocation [73], sound transmition [74], or feeding apparatus [75]. On the other hand, there is little facial soft tissues surrounding the skull of modern aquatic or semi-aquatic reptiles [76, 77]. Despite our general lack of understanding in the auditory mechanisms or predatory behavior of plesiosaurs, it is assumed here that they possessed a craniofacial musculature arrangement similar to those in modern aquatic reptiles. It has been demonstrated that the limited amount of craniofacial soft tissues of leatherback turtle (*Dermochelys coriacea*) are sufficient to maintain body heat in cold environments [78], and an analogous mechanism might also be present in plesiosaurs.

The fossil of *M. fernandezi* indicates that the necks of plesiosaurs were thicker *in vivo* than traditionally reconstructed [46]. In both extant aquatic mammals and reptiles, the neck outline forms a smooth curve connecting the skull and the trunk. A thick neck may exhibit a one-to-many mapping between form and functions. It may effectively prevent heat loss in high latitude regions or deepwater environments, or provide hydrodynamic advantages, thereby compensating for the energy cost required by nurturing the tissues [38]. Large amounts of muscles are present in the necks and temporal regions of crocodiles, enabling them to produce great force during predation [79]. Some thalassophonean pliosaurs were also macrophagous predators, evidenced by both biomechanical analyses [80] and the stomach contents [34], thus they might also possess well-developed neck muscles as in crocodiles.

Previous studies seldom discussed the amount of soft tissues around the ribcages of plesiosaurs, primarily due to the very recent discovery of *M. fernandezi*. In most previous publications, the body outlines were not provided or drawn close to the skeletons (e.g., [81, 32, 40]). The only study that quantitatively restored the soft tissues in the trunk region according to *M. fernandezi* is [36]. The width of soft tissue impressions in this specimen is 25% thicker than the ribcage ([46]: Fig. 3), which was employed as a criterion for soft tissue reconstrution in this study. However, it warrants caution that its ribcage has entirely collapsed during taphonomy, with the dorsal ribs detached from the transvere processes, and whether the ribs underwent caliper rotation relative to their life positions remains unclear. In other words, the width of ribcage in the current specimen may either overestimate or underestimate the original dimension *in vivo*. Furthermore, the soft tissue traces may also be thickened during taphonomy, introducing additional uncertainty in reconstruction. The 25% amount of soft tissue, which was applied as a reconstruction criterion, is derived directly from the specimen and works as a compromising option. In addition to the amount of soft tissues lateral to the ribcage, many aspects of soft tissue arrangement in the trunk region of plesiosaurs remain unclear (e.g., the amount of epaxial soft tissues). This leaves room for inference, which can be based on plesiosaur physiology and behavior, muscular restoration and comparison with modern analogues.

Despite their taxonomic position, plesiosaurs were physiologically similar to aquatic mammals, exemplified by endothermy and high metabolic rate (within the range of avians [82]), high body temperature (33 °C to 37 °C [83]), and viviparity [84, 85]. Modern marine mammals like cetaceans and pinnipeds possess thick blubber beneath the skin, which plays the roles of thermal barrier, energy storage, hydrodynamic streamlining and buoyancy regulation [86, 87, 88, 89]. Plesiosaurs, on the other hand, seemed to prefer high latitude and cold water environments. Many species are known to inhabit in or seasonally migrate to polar regions [90, 72, 91]. Without fur or hair, plesiosaurs were also likely to possess thick fat tissue to prevent heat loss. This is supported by the abundant soft tissues in *M. fernandezi*, which was, however, a species dwelling in warm water environment [46, 92]. Cetaceans from high latitude regions generally tend to possess thicker blubber than those from warmer seas [93, 94]. It is possible that plesiosaurs also follow this rule, but a quantitative study is not applicable in the current stage.

The amount of soft tissues at the dorsalventral sides of the ribcage of a plesiosaur also needs to be restored according to modern analogues. The ribcages of extant aquatic tetrapods are surrounded by muscles and ligaments (known as “core” in relevant studies, see [95] for example), outside of which are fat (blubber) and skin. The thoracic cross-sections in aquatic mammals are usually consistent in shape to the corresponding core profiles formed by muscles and the ribcage [96, 97, 98]. Assuming the same design, the cross-sectional profiles of plesiosaurs were modeled as an enlarged version of the corresponding core. The restoration of the later, on the other hand, requires the incorporation of musculoskeletal information of plesiosaurs. In the dorsal vertebrae of some plesiosaurs, the neural spines project high above the transverse processes (and in turn, the dorsal ribs) [54, 45], so that a cone-like cross-section will be produced if connecting the top of the spine and the outer margin of the rib. However, restoration in this way may neglect the presence of epaxial muslces. In cetaceans that also possess tall neural arches, the epaxial swimming muscle *m. longissimus dorsi* runs along the postcranial skeleton and surrounds the neural arches, creating a well-rounded cross-sectional profile [99]. Some major elevator muscles of the limbs of plesiosaurs were also inserted to the neural arches and extended along the vertebral column [100, 101], thereby potentially creating a rounded body outline as well (Fig. 2G).

It is also noteworthy that sea turtles, penguins, and otariids, as several major groups of extant underwater flyers, may not be the ideal references for the epaxial outline of plesiosaurs due to the significant differences in muscular distribution or limb kinematics among these groups. The major forelimb muscles for locomotion in sea turtles gather at their pectoral girdles and plastrons without attachment to the vertebrae [77]. In penguins, both the elevator and depressor muscles of their wings are attached to the ventral side of the pectoral girdle, similar to those of flying avians [102]. Such a pattern of muscular distribution creates a exceptionally robust chest, a feature that was likely absent in plesiosaurs. The extension ranges of the glenohumeral joints in otariids are limited, leading to asymmetric strokes during swimming: the downstroke is propulsive, while the upstroke is a passive recovery [103, 104]. Therefore the stroke style applied by plesiosaurs, which predominantly consisted of dorsalventral motions [101], was quite different from that applied by otariids.

Contour fat was present in the tail region of *M. fernandezi* in life, creating a streamlined body outline from the trunk to the tail tip [46]. The caudal regions of some other plesiosaurs indicate greater tail flexibility than those of polycotylids, revealing their potential participation in propulsion or manoeuvre [105, 106]. A pygostyle-like structure form by the fused terminal caudals is phylogeneticly widespread among plesiosaurs (see [107] and citations therein). Previous studies interpreted this structure as the potential evidence for the presence of a tail fin, but the orientation (horizontal vs vertical), shape, and relative size of the fin remain unclear to date [108, 109, 110, 111, 49]. Consequently, the tail fins of the plesiosaur models were not reconstruted in this study, and ommiting them should have negligible impact on the total body volumes.

### Cautionary notes on the volumetric equations

Major limiations of the hybrid approaches include the inadequate coverage of taxonomic range during sampling, the inconsistent standards for modelling in different studies, and the small sample size [14]. The methodology proposed in this study aims to mitigate the first two issues, but the last one remains inevitable since the number of models is strictly constrained by the fossil materials available. A rigorous skeletal reconstruction of plesiosaurs must be based on fossil specimens with at least one girdle elements exposed to determine the width and height of the ribcage. Such a threshold precludes the modelling of multiple well-preserved fossils (e.g., *Rhomaleosaurus cramptoni*; *Hauffiosaurus tomistomimus*; *Archaeonectrus rostratus*;), since their girdle elements are entirely obscured by the overlying ribs. Moreover, anatomical information including the intercervical distance, rib slant angles, spinal curvature is not always available due to fossil incomepleteness, necessitating the presence of both functionally analogous and phylogenetically related taxa as reference. When such referential taxa are absent, the modelling attempt is also precluded (e.g., *Brancasaurus brancai*; *Hauffiosaurus zanoni*). The current sample set contains 6 elasmosaurids, 2 cryptoclidids, 4 polycotylids, 8 pliosaurids, 2 rhomaleosaurids, and 2 microcleidids, covering all but one major clades of plesiosaurs. The only exception is Leptocleididae, which includes only a handful of taxa and is generally known from fragmentary fossils. The holotype of *Nichollssaura borealis* is an articulated and almost complete specimen, but the girdle elements are obscured [39]. However, this clade is phylogenetically bracketed by Polycotylidae and Cryptoclididae [55, 56], both of which are sampled in this study.

The current dataset contains plesiosaur models with various body plans, including the two taxa known to possess the highest and lowest number of cervical vertebrae respectively (75 in *Albertonectes vanderveldei* [42]; *>*8 [estimated at 11] in *“Monquirasaurus” boyacensis* [112]). Consequently, trunk length and DDV should be robust proxies of body volume against the votatile body proportions of plesiosaurs. Previous studies have demonstrated the limited validity of regression models when predicting the sizes of individuals which fall outside the range of the samples [18]. From the modest *Mauriciosaurus fernandezi* (which is an osteological immature individual [46]) to the gigantic *Sachicasaurus vitae*, the plesiosaur models created in this study range from 0.0776 m^3^ to 12.2288 m^3^ (Fig. 3AC), covering sufficient breadth to accommodate the size diversity across most plesiosaurs. It is also noteworthy that *M. fernandezi* is not included in the dataset of DDV-volume equation, since its dorsal vertebrae are obscured by the girdle elements. The smallest sample in that model is *Seeleyosaurus guilelmiimperatoris* SMNS 12039, which is still smaller than most plesiosaurs in body size. On the other hand, fragmentary fossils of multiple pliosaur individuals from the late Jurassic of Europe, empirically referrable to *Pliosaurus* sp, indicate body sizes larger than that of *S. vitae*. Investigation of their body sizes not only delineates the applicable boundaries of the volumetric equations, but also allows exploration for the potential maximum body size that was reached by plesiosaurs. The fossil materials of multiple pliosaur individuals indicate that they possibly reached or exceeded 20 metric tons in life (see the Supplementary Text for a brief body size revision of the mandible OUMNH PAL-J.010454), therefore caution is recommended when extrapolating the volumetric equations to them.

### Applicability and limitations of the paradigm

Most amniotes employ costal aspiration to ventilate their lungs, with the only extant exceptions being turtles [113]. The rib kinematics has been frequently investigated in physiological studies [114, 29], but was rarely discussed in skeletal reconstructions. The dorsal ribs are not properly articulated to the vertebral column in many mounted specimens, hence volumetric data derively directly from them should be treated with caution (pers. orbs. on mounted specimens during visits or through 3D scanns). In 3D modelling, multiple rotational and translational adjustments of the ribs are typically required to achieve proper positioning. Such complex manipulations can’t be implemented in 2D environments, since the ribs and vertebrae are all planar images. The lack of commutativity in 3D rotation contributes to the uncertainty in reconstructing the rib orientations. However, a decrease in the width of ribcage is always, at least partially, compensated by the increase in depth. The sensitivity tests performed in this study also imply that such uncertainty induces acceptable amount of discrepancy in volumetric estimates, at least in plesiosaurs (difference < 3.5% in all models). On the other hand, the current theoretical framework does not incorporate an angle to quantify the rib adduction (i.e., the extent of caliper rotation), which is determined during the alignment of vertically oriented rib planes with the vertebral images, prior to angle *θ*_1_ rotation. For bicapitate taxa, the the angle can be established according to the spatial positions of the diapophysis and parapophysis, whereas greater uncertainty may exist in clades with unicapitate ribs. But such uncertainty would result in minimal impact on the cross-sectional area, analogous to the effects observed when altering rotational sequences. When integrating human errors in physical model assembly and 3D modelling, the uncertainty in rib orientation appears to be an unavoidable issue of all VD approaches. Nevertheless, this study recommends future researchers to incorporate the knowledge of costovertebral joints and conduct inference of a static version of rib orientations.

This study proposes that prioritizing rotation angle *θ*_1_ rather than *θ*_2_ offers methodological advantages. In amniotes with bicapitate ribs, the line connecting the diapophysis and parapophysis forms an axis for rib rotation during costal aspiration [28]. For clades with single-headed ribs, the elliptical diapophyses in certain taxa serve a similar function [113]. Considering angle *θ*_1_ first essentially determines the orientation of this axis, enabling angle *θ*_2_ rotation to simulate the dynamic pathway of rib rotation during respiration. It is noteworthy that angle *θ*_1_ may exibit negative values in some clades (i.e., the rib inclines forward in lateral view if *θ*_2_ = 0°, exemplified by theropod dinosaurs [27]). Nevertheless, the mathematical formulae developed here remians valid due to the parity properties of trigonometric functions. In practice, the *θ*_1_ → *θ*_2_ sequence facilitates more efficient implementation of the CSM protocol used in this study, which defines the maximum cross-sectional height as the identidy segment, than *θ*_2_ → *θ*_1_. The cross-sectional profile is created by connecting the truncated ribs, of which the projected widths need to be calculated. For the *θ*_1_ → *θ*_2_ sequence, the projected width of each truncated rib 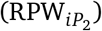 is given by RPW_*iP*_ = RPW_*i*_ *·* cos *θ*_2_, while it is RPW_*iP*_ = RPW_*i*_ *·* cos *θ*_2_ −RPH_*i*_ *·* sin *θ*_1_ *·* sin *θ*_2_ for *θ*_2_ →*θ*_1_ sequence (see the Supplementary Text for the deduction process). Therefore the former requires less sophisticated calculation during modelling.

As with all VD approaches, the programme of ribcage reconstruction followed by the CSM proposed in this study incorporates several assumptions that may not hold true across all amniotes. It assumes that the ribcage reliably reflects the soft thoracic outline, which is agnostic in most extinct taxa, and it’s inapplicable to armored taxa either (e.g., turtles and placodonts). The rib rotation protocol presumes that the orientation can be inferred from vertebral morphology, but may encounter limitations for unicapitate ribs with hemispherical costovertebral articulations (e.g., present in some squamates [113]). Furthermore, the dorsal migration of parapophyses in some clades (e.g., some mammals [115], and the posterior dorsal vertebrae of sauropod dinosaurs [116]) causes the capitulum to deviate from the plane defined by the tuberculum and rib shaft, thereby violating the assumption of rib simplification as a 2D structure. The axis formed by the diapophysis and parapophysis in these taxa theoretically constrains the range of both pump-handle and bucket-handle rotations, although the range of motion doesn’t entirely hinges on the morphology [28]. In such cases, the morphology of the vertebrae should to be taken into consideration to determine angles *θ*_1_ and *θ*_2_, and extra inspections are required to ensure the capitulum remains articulated to the parapophysis during rib rotation.

While subjectivity of the reconstruction process remains inevitable, the paradigm proposed in this study quantifies the relative sizes of skeletal elements, and facilitates the volumetric estimation within a specific clade even when only fragmentary fossil materials are available. This study provides formulae to estimate the body volumes of plesiosaurs for the first time, thereby partially filling the gap in estimating the body masses of secondarily aquatic tetrapods caused by inapplicability of existing ES approaches (e.g. the equation of terrestrial tetrapods based on stylopodial circumference [15, 16]). However, some limitations of the VD approaches are inherited. The time investment in modelling is enlarged during the exploration for a uniform set of reconstruction criteria. If the established criteria were realized inapplicable to some taxa, all previous models need to be dismantled and assembled again. Furthermore, the principle of VD approaches, which is utilized to generate samples for regression models, hampers the convenient test for uncertainty in skeletal dimensions (e.g. rib length).

### Enlightenment for future studies

As shown in Table 1, some large elasmosaurids attained body lengths comparable to or larger than those of giant thalassophonean pliosaurs, but their estimated volumes are substantially smaller, due to the long neck that occupies more than 50% of body length in multiple taxa. The maximum body volume of plesiosaurs was thus greatly reduced subsequent to the extinction of pliosaurs in Turonian [117]. On the other hand, the size evolution of Cryptoclidia (Fig. 3C) exhibits a deaccelerate pattern compared with other plesiosaur clades. The potential shifts in evolutionary dynamics of body size and the integrated variation in body plans remains a promising research scope for the future. A summary of size evolution within each plesiosaur clade was included in a previously preprinted version of this manuscript but not presented here due to consideration for article focus. This paper offers tools for quick and convenient volumetric estimation of various plesiosaur taxa, thereby facilitating the determination of the mode and tempo of plesiosaur size evolution. In addition, the phylogenetic analysis based on Bayesian inference illuminates the diversification and extinct rates of plesiosaurs within each time bin, which deserve futhur interpretation. These topics will be further investigated and explored in future publications of the current author (in prep.).

## Methods

### Phylogenetic data and analysis

The phylogeny of plesiosaurs is in a state of flux. Although research over the past decade has established consensus regarding the phylogenetic relationships of major clades, the internal topology within each lineage remains poorly resolved [118, 119, 120]. Almost all current phylogenetic analyses of plesiosaurs are based on modified versions of the character matrix constructed by [55]. Researchers have added different operational taxonomic units (OTUs; in this case, species) and morphological characters to the original matrix, resulting in incompatibilities between them. Furthermore, all existing character matrices lack some species included in this study, necessitating a reorganization and consolidation of them.

A recent version of the character matrix was selected from [121], and substantial modifications were made to it using Mesquite 3.8.1 [122]. The missing OTUs were obtained from other matrices, with their scores revised. The modification to the names of some polycotylids by [107] was also adopted here. For the unique characters in this matrix, the newly added OTUs were scored based on published literature, while undescribed or uncertain features were assigned a “?”, pending a thorough reappraisal of these specimens, following [120]. A paper that revised the scores of *Seeleyosaurus guilelmiimperatoris* was published during the preparation of this study, revealing a slightly different topology within Microcleididae [49]. Their modification wasn’t incorporated into the current matrix, since they announced plans for a comprehensive reassessment of the genus *Microcleidus* in the near future. The resulting matrix contains 290 characters and 146 OTUs, which is one of the largest plesiosaur matrix to date. A Bayesian method, the skyline variant fossilzed birth-death model (SFBD) model, was used to investigate the the topology of plesiosaur phylogeny. It can jointly estimate tree topology, divergence times, and other macroevolutionary parameters across different time bins [123, 124]. To investigate potential heterogeneity in the phenotypic evolution rates of plesiosaurs, the constant rate assumption was relaxed and an uncorrelated lognormal clock was used to estimate the branch-specific rates [125]. Given the complexity of the results and the focus of this study, the construction of character matrix and the phylogenetic analysis will be elucidated in detail in another recent publication of the current author (in prep.).

### Regression

The selected variables (see Fig. S2 for measuring criteria) were first log_10_-transformed prior to the model fitting. The OLS models were fitted using the lm() function implemented in base R stats package version 4.3.3 [126], and the PGLS models were fitted with the gls() function from the nlme package version 3.1-164 [127]. Considering that small sample sizes may negatively affect the estimation of phylogenetic signals, the corBrownian autocorrelation structure rather than the corPagel or corBlomberg was employed to accommodate phylogenetic non-independence. For the skull-neck dataset, a nonlinear regression model was fitted by using the drm() function implemented in the drc package version 3.0-1 [128], with a four-parameter log-logistic function activated (fct = LL.4()). The performance of the regression models was compared using the AICc and 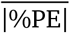 values. The AICc value quantifes the fitness of model to data, while the 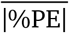 value evaluates the performance in prediction [57, 14]. To compute the later, each specimen was iteratively removed from the dataset as the testing set, with the remaining specimens serving as the training set for the regression modelling. The resulting model was then used to predict the value of the testing set, and the antilog-scaled prediction value was compared with the observed value:

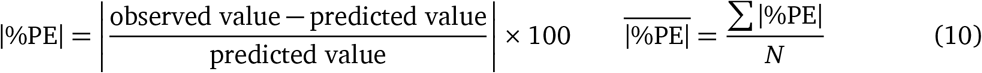

### Pleiosaur reconstruction

A general outline of 2D skeletal reconstruction of extinct amniotes is provided in the Results section, and some supplementary details of its application in plesiosaurs are provided here. All models were constructed in AutoCAD, with the software resolution set to 0.0001 mm. Measurements were cited from literature, or taken from strict dorsal or lateral view photos from either publications or colleagues. The rib rotation protocol required to reconstruct ribcage cross-sections was implemented using the Rotation and Rotation3D commands in AutoCAD. All models were created under the *θ*_1_ → *θ*_2_ rib rotation sequence, while a *θ*_2_ → *θ*_1_ version was also created for the sensitivity test. The difference of the two versions were calculated as 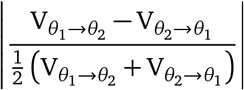

### Main axis

The criteria for calculating spinal length are described first since estimating the SKL of a plesiosaur requires the neck length and the cervical number. If the selected specimen is articulated or broken into several blocks, the spinal length was acquired by summing the measurements along the column. However, the fragmentary and disarticulated nature of many plesiosaur fossils necessitates the estimation of their spinal lengths. In the case of relatively complete vertebral columns, where only a small number of vertebrae are absent or unmeasurable, the lengths of these vertebrae were restored by calculating the mean value of the adjacent ones. Should multiple vertebrae be absent or unmeasurable, an individual with a complete vertebral column from a closely related or anatomically analogous species (e.g. one that possesses the same number of cervical vertebrae) was selected for reference.

The dimensions of the intercervical cartilage exhibit variation across different plesiosaur species, with a negative correlation with the cervical number. As argued in [129], the increased intercervical distance in short-necked taxa might play a compensatory role to increase the neck flexibility. The relative amount of intercervical distances of different plesiosaur clades were summarized from fossil specimens with an articulated neck, and employed as references to reconstruct other plesiosaurs (13% for rhomaleosaurids, as in *Rhomaleosaurus thorntoni* [130]; 25% for thalassophonean pliosaurs, as in *Sachicasaurus vitae* [131] and *Brachauchenius lucasi* [34]; 8% for elasmosaurids, as in *Libonectes morgani* [132]; 9% for late Jurassic and early Cretaceous cryptoclidids, as in *Ophthalmothule cryostea* [50]; 13% for polycotylids, as in *Mauriciosaurus fernandezi* [46]). On the other hand, the dimensions of intervertebral cartilage from the pectoral, dorsal, and sacral regions appear to be approximately 10% of the vertebral length, as evidenced by multiple articulated plesiosaur specimens (e.g., *B. lucasi* [34], *R. thorntoni* [130], *Elasmosaurus platyurus* [42]). The estimation of intercaucal distance is not considered warranted here, since most plesiosaur specimens with a disarticulated tail do not contain a complete caudal series. Hence the trunk-tail equation was employed to predict tail length in this study. For specimens without a preserved skull, the SKL was predicted using the skull-neck equation. Another estimate was generated based on the ratio of SKL to neck length in a congeneric species if feasible, and the final SKL was set as the mean of both estimates. The basal cranial length (BSL [34]; Fig. S2), measured from the snout tip to the occipital condyle, was then computed using the SKL/BSL proportion of a close relative.

### Ribcage and body shape

To reconstruct the body shape of a plesiosaur, the skull and vertebral column were created first. The curvature of the spine exerts a substantial influence on plesiosaur body shape and hydrodynamic properties, thus it needs to be carefully inferred. Some plesiosaur fossils indicate that the anterior end of scapulae is aligned with the first pectoral vertebra, and the acetabulum is aligned with the second sacral vertebra (e.g. *“Monquirasaurus” boyacensis* [112], *Albertonectes vanderveldei* [41]). The trunk length was drawn as a horizontal line segment, and another curve was created to connect the two endpoints of the segment. The length of the curve was stretched to approximate the calculated length along the vertebral column (pectoral+dorsal+sacral 1), and the vertebrae were then placed along the curve.

To enable the calculation of body volume and surface area, three thoracic cross-sections were reconstructed for each plesiosaur model: the glenoid cross-section (the vertical plane containing the pectoral glenoid), the acetabulum cross-section (the vertical plane containing the pelvic acetabulum), and the middle cross-section (at the middle of the other two cross-sections). Each cross-section was split into the dorsal part and the ventral part. All the dorsal parts of the three cross-sections include vertebrae and ribs. In the glenoid and acetabulum cross-sections, the ventral part consists of the coracoids and pubes respectively, and the ventral part of maximum cross-section is gastralia. The size and shape of each cross-section are determined in this stage.

The dorsal aspects of the three cross-sections were reconstructed by identifying the rib orientation. All dorsal ribs in plesiosaurs are single-headed, attached to the transverse processes of the neural arches. The sizes of angles *θ*_1_ and *θ*_2_ vary greatly across plesiosaur species (e.g., max angle *θ*_1_ = 40° in *Cryptoclidus eurymerus* [51], and 30° in *Hydrotherosaurus alexandrae* [43]). Judging from well-preserved plesiosaur fossils, the sizes and slant angles of transverse processes shift gradually. The transverse processes often increase in size till middle of the dorsal region and become shortened afterwards. Angle *θ*_1_ also increases gradually from 0° in anterior dorsals to maximum in middle of the dorsal region, and keeps almost constant in the posterior half of dorsal series but decreases to 0° again rapidly when approaching the sacral region. On the other hand, angle *θ*_2_ increases monotonically from 0° to maximum, and remains constant with no recovery. Such an arrangement of rib orientation, which seems to be widespread among plesiosaurs, was described by previous researchers or can be observed from multiple well-preserved fossils ([54, 45, 133]; pers. orbs on *Avalonnectes arturi*). The ribs also change in shape and size throughout the dorsal region. They typically increase in length gradually in anterior half of the dorsal region, then become shorter and straighter in the posterior half [134, 54]. The dorsal ribs of some plesiosaur specimens are not preserved or can’t be measured directly because of their mounted state. In these circumstances, the maximum arc length of the dorsal ribs was estimated using the trunk-rib equation.

As mentioned above, the pair of dorsal ribs projected to the glenoid cross-section was placed with angles *θ*_1_ and *θ*_2_ both set to 0, and only angle *θ*_3_ was considered. For specimens with uncertain rib lengths, the arc length of the rib projected to the glenoid cross-secion was reconstructed using body proportions of closely related species (arc length at the glenoid level/maximum rib arc length; 60% for rhomaleosaurids, as in *Rhomaleosaurus cramptoni* [135]; 80% for thalassophonean pliosaurs, as in *“Monquirasaurus” boyacensis* [112]; 66% for elasmosaurids, as in *Wapuskanectes betsynichollsae* [39]; 70% for polycotylids, as in *Dolichorhynchops osborni* [136]; 68% for cryptoclidids, as in *Cryptoclidus eurymerus* (pers. orbs from open source 3D scann from https://sketchfab.com/bcdh); 72% for microcleidids, as in *Microcleidus homalospondylus* [105]). On the other hand, the height and width of the dorsal part of the acetabulum cross-section can’t be determined in this way because of the shrunken and straightened ribs. It is assumed here that this cross-section has the same width as that of the glenoid region, following some articulated or mounted plesiosaur fossils (e.g., *Mauriciosaurus fernandezi* [46]; *Meyerasaurus victor* [137]; pers. orbs on the ventral view photos of *Thalassomedon hamingtoni*). The dorsal outline of this cross-section was constructed by scaling and stretching the glenoid cross-section after its width and height were determined.

The coracoids, pubes and ischia in plesiosaurs are plate of bones, and the two symmetrical components form an obtuse angle at the midline junction of the body, creating a “V”-shape from the main side view [54]. The size and orientation of the dorsal ribs determine the width of the trunk, and a mathematical method was used to reconstruct the angle of intersection between the coracoids or the pubes. Given the widths of the girdle elements at the glenoid and acetabulum, the heights of the glenoid and acetabulum cross-sections were calculated using the Pythagorean theorem. Height of the ventral part of the maximum cross-section was obtained with the following approach: average height of the ventral parts of the glenoid and the acetabulum cross-sections was taken, then the result was scaled by multiplying the inverse of rib length distributional proportion which was used in estimating rib lengths at the glenoid level.

### Flipper and soft tissue

The four limbs of all plesiosaurs were hydrofoil-like flippers that exhibited hyperphalangy [138, 139], and their distal ends were incomplete in most fossil specimens. Therefore, restoration is frequently required. If the propodial of the flipper under study is preserved, total length of the whole limb was first estimated from propodial chord using the equation provided in [140]. Then the missing elements were restored according to the shapes and proportions of those from a close relative. If a pair of forelimbs or hindlimbs are entirely missing, flipper lengths were restored by comparison with congeneric or kin species. The cross-sections of all flippers were approximated using a same hydrofoil profile, which was employed in a previous study investigating plesiosaur locomotion [141]. The rationale for the soft tissue reconstruction along the main body axis is elucidated in Discussion, and the criteria are briefly reiterated here. Barely any craniofacial soft tissues were added to the plesiosaur models, and the three cross-sections in the trunk region were enlarged by 25%. Five percent extra tail length representing soft tissues were added to the posterior end of each model, following the soft tissue traces in *Seeleyosaurus guilelmiimperatoris* MB.R.1992 [49]. The lack of complete soft tissue traces revealing the profile of flippers *in vivo* causes a dilemma to standardized reconstruction. In order to quantitatively restore the flippers of all plesiosaur models under the same criteria, the reconstruction procotol proposed in [141] was employed in this study.

### Volumetric computation

The CSM was employed to compute the body volume of each plesiosaur model [24]. Three crosssections for the ribcage has been established above, and an ellipse at the quadrate level was introduced to approximate the cross-sectional profile of the skull. The major axis and minor axis of the ellipse was defined as the skull width and skull height at the quadrate level (estimated from a close relative if the skull is missing). The main body axis is thus partitioned into 5 slabs by the four cross-sections (Fig. 1). A constant shape was assumed for the slab at each end of the body (i.e., the first one and the last one), while others were assumed to possess gradually changing cross-sections. Each limb was treated as a single slab, and the cross-section of a hydrofoil used in [142] was employed here. Each slab was sliced into 100 subslabs, which far exceeds the number required by stable estimation of the CSM [24]. The model slicing was performed in AutoCAD, and the computation was performed in Excel.

### Mode and tempo of plesiosaur size evolution

The body volumes of 113 plesiosaur taxa were estimated based on the model, trunk length, DDV, ischium length, femur chord, and SKL×CN. The references for data source are available as a pdf file in the Supplementary Material. To investigate the rate heterogeneity in the evolution of plesiosaur body size, a variable-rates model implemented in BayesTraits 4.0.0 [143] was fitted to the log_10_-transformed data and the pruned MCCT. Rate shift was evaluated using a reversible jump Markov Chain Monte Carlo algorithm (rjMCMC). The tree was rescaled using the Pagel’s *λ* prior to the iterations. The analysis takes 22000000 iterations, with the first 2000000 set as burn-in. The results were sampled every 20000 iterations, then the mean rate of each branch was summarized and mapped to the pruned MCCT. To mitigate potential bias caused by potentially unreliable size proxies and osteologically immature individuals, a second analysis was performed based on the reduced dataset consisting of body volumes of mature individuals, estimated from the models, trunk length, or DDV. To futher account for the impact from unstable phylogeny, the second analysis on the reduced dataset was replicated on 100 trees randomly selected from the post burn-in samples. A homogeneous Brownian model was also fitted, and its performance compared with the heterogeneous model was evaluated using log Bayes Factors, which was calculated from marginal likelihoods of the two models obtained from the stepping-stone sampling procotol (100 stones per run for 1000 iterations). Convergence of the parameters was checked by examining the ESS values (ESS*>*200) using the R package coda version 0.19-4.1 [144]. Tree manipulation was carried out using the R package ape version 5.8 [145]. Visualization was implemented using the packages ggplot2 version 3.5.1 [146], ggforce version 0.4.2 [147], ggtree version 3.13.1 [148], deeptime version 2.0.0 [149], and cowplot version 1.1.3 [150].

## Supporting information

References for data

skull-neck data

proxy for volume

raw body volume data

trunk-rib data

trunk-tail data

## Acknowledgements

This paper presents part of my investigation results on plesiosaurs during the past four years. I warmly thank Benedon Paratodus (pseudonym, used as requested), Frank Fang, Emo Young, Shinya Noguchi, Frederick Dakota, Yang Song, Y.-W. Fang, Devin LYu, Lingcheng Liu and Andy Thomas for discussion and their constant support during the preparation of this study. I also thank Espen Knutsen, Richard Forrest and Leslie Noè, who offered comments and suggestions on the plesiosaur reconstructions. Nikolay Zverkov, Jørn Hurum, Nigel Larkin, Luis Spalletti, Zulma Gasparini, Anna Krahl, Carla Crook, Eric Buffetaut, Peggy Vincent, Glenn Storrs, Bruce Schumacher and Eberhard Frey are thanked for kindly sharing their knowledge or providing photos.

## Notes

### Competing Interest Statement

The authors have declared no competing interest.

### Summary of Updates

The theoretical framework and all existing skeletal reconstructions modified and improved; several new models added; manuscript partially rewritten for clearer elaboration.

