## Supplementary Text_Figures_Tables for "Body reconstruction and size estimation of plesiosaurs: enlightenment on the ribcage restoration of extinct amniotes in 2D environments"

Ruizhe Jackevan Zhao

### Supplementary Text

#### Rib rotation $\theta_2 \to \theta_1$

Angle $\theta_2$ is considered first, and the rib plane is rotated around the AD axis by $\theta_2$ degrees. Then RPH projected to the vertical plane in this moment ($\mathrm{RPH}_{P_1}$) equals RPH, while the projected RPW ($\mathrm{RPW}_{P_1}$) equals RPW$\cdot \cos\theta_2$. Then the rib plane is flipped around the AB axis by $\theta_1$ degrees, and the projected height ($\mathrm{RPH}_{P_2}$) and width ($\mathrm{RPW}_{P_2}$) are given by:

$$\mathrm{RPH}_{P_2} = \mathrm{RPH} \cdot \cos\theta_1 \qquad \mathrm{RPW}_{P_2} = \mathrm{RPW}_{P_1} - \mathrm{RPH} \cdot \sin\theta_1 \cdot \sin\theta_2 \tag{1}$$

After the determination of rib orientation inferred from the vertebral morphology, the costovertebral system is rotated around the mediolateral axis by $\theta_3$ degrees. Then the projected height and width are $\mathrm{RPH}_{P_3} = \mathrm{RPH}_{P_2} \cdot \cos\theta_3$ and $\mathrm{RPW}_{P_3} = \mathrm{RPW}_{P_2}$, respectively. Subsequent to the reconstruction of the ribcage in lateral view, a specific vertical plane is drawn to truncate multiple dorsal ribs. The truncated height and width of each rib plane ($\mathrm{RPH}_i$ and $\mathrm{RPW}_i$, $i = 1, 2, \cdots$) is then determined, the projected width ($\mathrm{RPW}_{iP}$) equals $\mathrm{RPW}_i \cdot \cos\theta_2 - \mathrm{RPH}_i \cdot \sin\theta_1 \cdot \sin\theta_2$,

#### On some giant Jurassic pliosaurs

OUMNH PAL-J.010454, a mandible that was classified to *Stretosaurus* [1], *Liopleurodon* [2] and *Pliosaurus* [3] before, was restored as 2875 mm in length [1]. Length of the imperfect mandible before restoration was 7 feet (about 2134 mm), as briefly mentioned in [4]. The mandible is currently displayed in a glass exhibition case, precluding first-hand measurements, hence the photogrammetric method was used here to investigate its size. There exists a breakage behind the dentary on each ramus of the reconstructed mandible, and length of the mandible anterior to breakage matches this value (pers. orbs.). It was argued in [1] that "...the posterior part of the left ramus has come to light... the total length would have been more than 3000 mm". There are indeed two associated lower jaw fragments from a single individual (OUMNH PAL-J.050376 and OUMNH PAL-J.050377) discovered in the same pit with OUMNH PAL-J.010454 and they match in size [3]. It is not certain which specimen was referred to in [1], but if it was OUMNH PAL-J.050376, the anteroposterior length of the mandible should be around 2.6 m. Assuming an identical body proportion shared with *Pliosaurus* cf. *kevani*, its body mass might reach 20 metric tons. It is not the only example indicating that Jurassic pliosaurs might reach 20 t in mass. Some giant cervical vertebrae described in [5] and an isolated dorsal rib that is 122 cm in chord length [6] all indicate a similar body size.

### Institutional abbreviations

**TMP**, Tyrrell Museum of Palaeontology, Drumheller, Alberta, Canada; **SGO.PV**, Museo Nacional de Historia Natural, Santiago, Chile; **UCMP**, Museum of Paleontology, University of California at Berkeley, Berkeley, California; **SDSM**, South Dakota School of Mines, Rapid City, U.S.A; **DMNH**, Denver Museum of Nature and Science, Denver, U.S.A.; **MLP**, Museo de La Plata, Buenos Aires, Argentina; **MChEIO**, Museum of Chuvash Natural Historical Society, Chuvashia, Russia; **NHMUK**, Natural History Museum, London, U.K.; **FHSM**, Fort Hays State University, Sternberg Museum of Natural History, U.S.A.; **KUVP**, Natural History Museum, University of Kansas, Lawrence, U.S.A.; **INAH**, Instituto Nacional de Antropología e Historia, Saltillo, Mexico; **YPM**, Yale Peabody Museum, New Haven, U.S.A.; **MCZ**, Museum of Comparative Zoology, Harvard University, U.S.A; **GPIT**, Geologisch-Paläontologisches Institut Tübingen, Tübingen, Germany; **MJACM**, Museo El Fósil, Vereda Monquirá, Colombia; **CAMSM**, Sedgwick Museum of Geology, Cambridge, U.K.; **PMO**, Palaeontology Museum, Natural History Museum, Oslo, Norway; **SMNS**, Staatliches Museum für Naturkunde, Stuttgart, Germany; **MMM**, Museé Municipal de Millau; **OUMNH**, Oxford University Museum of Natural History, Oxford, U.K.; **MB**, Naturkundemuseum Berlin, Berlin, Germany;

### Supplementary Tables

**Table S1. Parameters of phylogenetic generalized least squares (PGLS) models and their comparison with corresponding ordinary least squares (OLS) models.** All countinous measurements are in meters (m), and all proxies and the volumes were $\log_{10}$-transformed before analysis. Abbreviations: N, sample size; AICc, sample-size corrected Akaike Information Criterion; SKL, skull length; CN, cervical number; DDV, dimensions of dorsal vertebrae (mean length×mean width×mean height of the dorsal centra); $\Delta AICc = AICc_{PGLS} - AICc_{OLS}$. See Fig. S2 for measuring criteria.

| Proxy | $Slope_{PGLS}$ | $Intercept_{PGLS}$ | N | P-value | $AICc_{PGLS}$ | $AICc_{OLS}$ | $\Delta AICc$ |
|---|---|---|---|---|---|---|---|
| SKL×CN | 3.2033 | -3.7970 | 16 | <0.001 | 12.90 | 3.13 | 9.77 |
| trunk | 2.6239 | -0.4694 | 24 | <0.001 | -33.36 | -46.93 | 13.56 |
| DDV | 0.9563 | 3.2957 | 14 | <0.001 | -7.35 | -20.12 | 12.77 |
| humerus length | 2.6767 | 1.0140 | 23 | <0.001 | 5.43 | 13.06 | -7.63 |
| humerus chord | 2.6457 | 1.8835 | 23 | <0.001 | 18.48 | 14.11 | 4.37 |
| femur length | 2.7884 | 1.0090 | 21 | <0.001 | 7.01 | 16.02 | -9.01 |
| femur chord | 2.5553 | 1.8061 | 21 | <0.001 | 6.69 | 1.38 | 5.31 |
| coracoid length | 2.2152 | 0.6776 | 16 | <0.001 | 5.91 | 1.49 | 4.41 |
| coracoid width | 2.9254 | 1.6963 | 22 | <0.001 | -4.80 | -3.81 | -0.98 |
| pubis length | 2.0993 | 0.9572 | 17 | <0.001 | 19.97 | 7.06 | 12.91 |
| pubis width | 3.1010 | 1.4998 | 19 | <0.001 | -5.29 | -3.50 | -1.78 |
| ischium length | 1.9712 | 0.9111 | 19 | <0.001 | 15.43 | 13.32 | 2.11 |
| ischium width | 3.0810 | 1.8726 | 19 | <0.001 | -0.01 | -8.70 | 8.68 |

**Table S2. Results of the sensitivity tests performed on the plesiosaur models.** $V_{\theta_1\rightarrow\theta_2}$ refers the volume of the $\theta_1 \rightarrow \theta_2$ model, while $V_{\theta_2\rightarrow\theta_1}$ refers to the volume of the $\theta_2 \rightarrow \theta_1$ model (see the manuscript for definitions). Difference is defined as $\left|\frac{V_{\theta_1\rightarrow\theta_2} - V_{\theta_2\rightarrow\theta_1}}{\frac{1}{2}\left(V_{\theta_1\rightarrow\theta_2} + V_{\theta_2\rightarrow\theta_1}\right)}\right|$.

| Taxon | $V_{\theta_1\rightarrow\theta_2}$ ($m^3$) | $V_{\theta_2\rightarrow\theta_1}$ ($m^3$) | Difference |
|---|---|---|---|
| *Albertonectes vanderveldei* | 3.0098 | 2.9204 | 3.02% |
| *“Aristonectes” quiriquinensis* | 4.0562 | 4.0407 | 0.38% |
| *Hydrotherosaurus alexandrae* | 1.8277 | 1.7679 | 3.33% |
| *Styxosaurus* sp. | 2.9987 | 2.9109 | 2.97% |
| *Thalassomedon hamingtoni* | 6.6125 | 6.5479 | 0.98% |
| *Vegasaurus molyi* | 0.8172 | 0.8142 | 0.37% |
| *Abyssosaurus nataliae* | 1.2057 | 1.2062 | 0.04% |
| *Cryptoclidus eurymerus* | 0.3613 | 0.3639 | 0.70% |
| *Dolichorhynchops osborni* | 0.4137 | 0.4170 | 0.78% |
| *Martinectes bonneri* | 1.2330 | 1.2599 | 2.16% |
| *Mauriciosaurus fernandezi* | 0.0776 | 0.0757 | 2.40% |
| *Polycotylus latipinnis* | 1.0292 | 1.0347 | 0.54% |
| *Kronosaurus/Eiectus* | 11.0968 | 10.9679 | 1.17% |
| *Liopleurodon ferox* | 1.7482 | 1.7229 | 1.46% |
| *“Monquirasaurus” boyacensis* | 9.6561 | 9.6252 | 0.32% |
| *Peloneustes philarchus* | 1.0075 | 1.0068 | 0.07% |
| *Pliosaurus* cf. *kevani* | 10.4712 | 10.4172 | 0.52% |
| *Pliosaurus funkei* | 10.5937 | 10.5397 | 0.51% |
| *Sachicasaurus vitae* | 12.2288 | 12.0180 | 1.74% |
| *Stenorhynchosaurus* | 2.0928 | 2.0695 | 1.12% |
| *Macroplata tenuiceps* | 0.6703 | 0.6793 | 1.34% |
| *Meyerasaurus victor* | 0.3639 | 0.3702 | 1.72% |
| *Microcleidus tournemirensis* | 0.2173 | 0.2175 | 0.11% |
| *Seeleyosaurus guilelmiimperatoris* | 0.1940 | 0.1890 | 2.65% |
| mean | | | 1.27% |

### Supplementary Figures

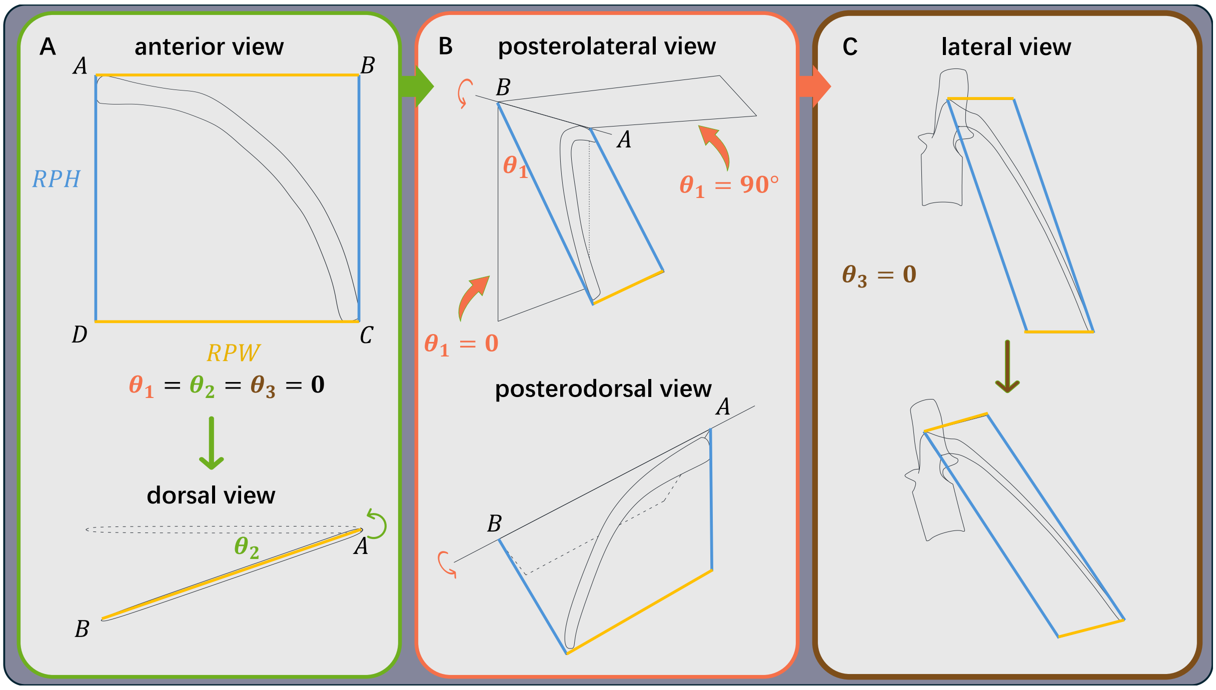


**Figure S1. The protocol of rib rotation ($\theta_2 \rightarrow \theta_1 \rightarrow \theta_3$).** (A) The vertically orientated rib plane ($\theta_1 = \theta_2 = \theta_3 = 0$) showing the rib plane height (RPH) and rib plane width (RPW), and lateral view of the rib plane that rotates around the AD axis by $\theta_2$ degrees. (B) Posterolateral and posterodorsal views of the rib plane that rotates around the AB axis by $\theta_1$ degrees. (C) Lateral view of the costovertebral system that rotates around the mediolateral axis by $\theta_3$ degrees.

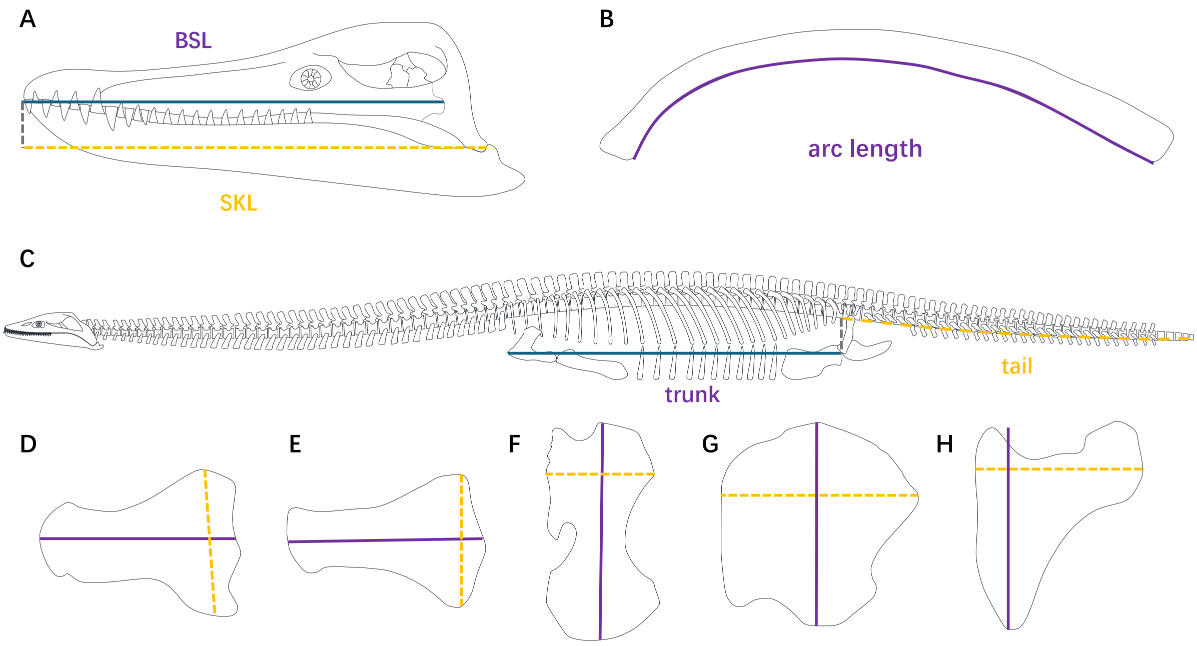


**Figure S2. Schematic images of the measuring criteria of skeletal elements.** (A) The basal cranial length (BSL, from snout tip to occipital condyle; solid line) and the skull length (SKL, from snout tip to quadrate; dashed line). (B) The arc length of a dorsal rib. (C) The trunk length (solid) and the tail length (dashed). (D) The humerus length (solid) and chord (dashed). (E) The femur length (solid) and chord (dashed). (F) The coracoid length (solid) and width (dashed). (G) The pubic length (solid) and width (dashed). (H) The ischial length (solid) and width (dashed).

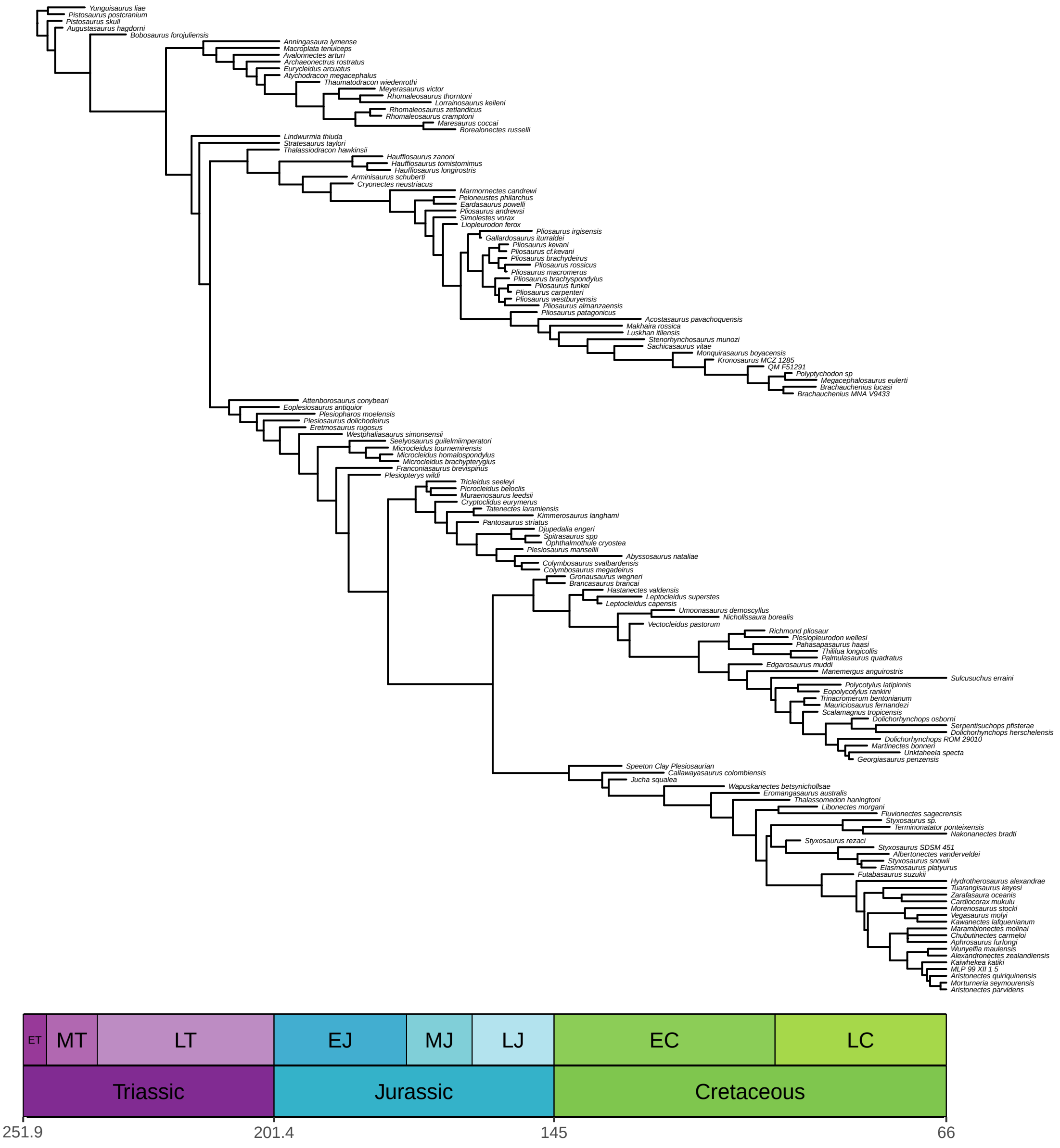


**Figure S3. The maximum clade credibility tree (MCCT) summarized from the post burn-in Bayesian tree samples.**

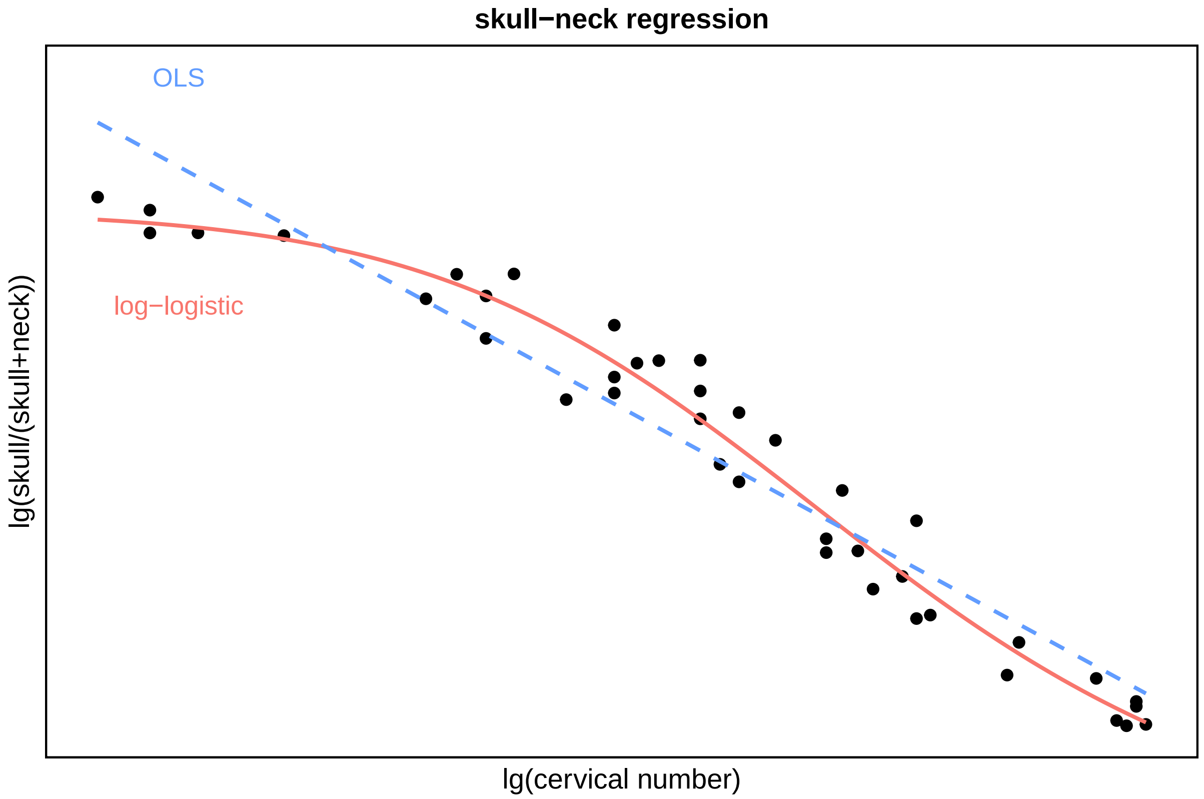


**Figure S4. Scatter plot and regression models of the skull-neck data set.** The blue dashed line represents the linear regression based on ordinary least squares (OLS), while the red curve represents the nonlinear regression based on a log-logistic function.
