## Supplementary material for "Body reconstruction and size estimation of plesiosaurs: enlightenment on the ribcage restoration of extinct amniotes in 2D environments": References for data

Ruizhe Jackevan Zhao

This file contains the references of the datasets used in this study.

### Neck-SKL regression.xlsx

| | |
|---|---|
| *Libonectes morgani* | Welles, 1949 [1]; |
| *Meyerasaurus victor* | Smith & Vincent, 2010 [2]; |
| *Sachicasaurus vitae* | Páramo-Fonseca et al, 2018 [3]; |
| *Kaiwhekea katiki* | Cruickshank & Fordyce, 2002 [4] |
| *Ophthalmothule cryostea* | Roberts er al, 2021 [5] |
| *Trinacromerum bentonianum* | Thurmond, 1968 [6] |
| *Thililua longicollis* | Bardet et al, 2003 [7] |
| *Hauffiosaurus tomistomimus* | Benson et al, 2011 [8] |
| *Styxosaurus SDSM 451* | Welles & Bump, 1949 [9] |
| *Dolichorhynchops osborni* | Carpenter, 1999 [10]; Bonner, 1964 [11] |
| *Hydrotherosaurus alexandrae* | Welles, 1943 [12] |
| *Thalassomedon haningtoni* | Welles, 1943 [12] |
| *Styxosaurus sp.* | Welles, 1952 [13] |
| *Edgarosaurus muddi* | Drunkenmiller, 2002 [14] |
| *Microcleidus homalospondylus* | Benson et al, 2012 [15] |
| *Plesiopterys wildi* | O'Keefe, 2004 [16] |
| *Plesiosaurus dolichodeirus* | O'Keefe, 2004 [16] |
| *Thalassiodracon hawkinsii* | O'Keefe, 2004 [16] |
| *Brancasaurus brancai* | Sachs et al, 2016 [17] |
| *Atychodracon megacephalus* | Smith, 2015 [18] |
| *Macroplata tenuiceps* | Ketchum & Smith, 2010 [19] |
| *Brachauchenius lucasi* | McHenry, 2009 [20] |
| *Kronosaurus queenslandicus* | McHenry, 2009 [20] |
| *Kronosaurus MCZ 1285* | McHenry, 2009 [20] |

| | |
|---|---|
| *Luskhan itilensis* | Fischer et al, 2023 [21] |
| *Attenborosaurus conybeari* | Benson et al, 2012 [15] |
| *Simolestes vorax* | O'Keefe, 2002 [22] |
| *Muraenosaurus leedsii* | O'Keefe, 2002 [22] |
| *Cryptoclidus eurymerus* | Brown, 1980 [23] |
| *Serpentisuchops pfisterae* | Persons et al, 2022 [24] |
| *Callawayasaurus colombiensis* | Welles, 1962 [25] |
| *Abyssosaurus nataliae* | Berezin, 2011 [26]; Berezin, 2019 [27] |
| *Nakonanectes bradti* | Serratos et al, 2017 [28] |
| *Rhomaleosaurus cramptoni* | O'Keefe, 2002 [22] |
| *Archaeonectrus rostratus* | Benson et al, 2012 [15] |
| *Microcleidus tournemirensis* | Bardet et al, 1999 [29] |
| *Thaumatodracon wiedenrothi* | Smith & Araújo, 2017 [30] |
| *Liopleurodon ferox* | Linder, 1913 [31] |
| *Peloneustes philarchus* | Linder, 1913 [31] |
| *Monquirasaurus boyacensis* | Noè & Gómez-Pérez, 2022 [32] |
| *Nichollssaura borealis* | Nagesan, 2017 [33] |

### Trunk-Rib regression.xlsx

| | |
|---|---|
| *Macroplata tenuiceps* | Ketchum & Smith, 2010 [19] |
| *Atychodracon megacephalus* | Smith, 2007 [34] |
| *Rhomaleosaurus cramptoni* | Smith, 2007 [34] |
| *Archaeonectrus rostratus* | Owen, 1865 [35] |
| *Peloneustes philarchus* | Linder, 1913 [31] |
| *Sachicasaurus vitae* | Páramo-Fonseca et al, 2018 [3] |
| *Stenorhynchosaurus munozi* | Páramo-Fonseca et al, 2016 [36] |
| *Monquirasaurus boyacensis* | Noè & Gómez-Pérez, 2022 [32] |
| *Liopleurodon ferox* | Vincent et al, 2024 [37] |
| *Plesiosaurus dolichodeirus* | Storrs, 1997 [38] |
| *Seelyosaurus guilelmiimperatori* | Dames, 1895 [39] |
| *Microcleidus homalospondylus* | Owen, 1865 [35] |
| *Cryptoclidus eurymerus* | Richards, 2011 [40] |
| *Abyssosaurus nataliae* | Berezin, 2019 [27] |
| *Tatenectes laramiensis* | O'Keefe et al, 2011 [41] |
| *Nichollssaura borealis* | Druckenmiller & Russell, 2008 [42] |
| *Dolichorhynchops osborni* | Willison, 1907 [43] |
| *Mauriciosaurus fernandezi* | Frey et al, 2017 [44] |
| *Wapuskanectes betsynichollsae* | Henderson, 2024 [45] |
| *Kaiwhekea katiki* | Cruickshank & Fordyce, 2002 [4] |

| | |
|---|---|
| *Albertonectes vanderveldei* | Henderson, 2024 [45] |
| *Vegasaurus molyi* | O'Gorman et al, 2015 [46] |
| *Hydrotherosaurus alexandrae* | Welles, 1943 [12] |

### Trunk-Tail regression.xlsx

| | |
|---|---|
| *Albertonectes vanderveldei* | Kubo et al, 2012 [47] |
| *Macroplata tenuiceps* | Ketchum & Smith, 2010 [19] |
| *Rhomaleosaurus cramptoni* | Smith, 2007 [34] |
| *Archaeonectrus rostratus* | Owen, 1865 [35] |
| *Plesiosaurus dolichodeirus* | Storrs, 1997 [38] |
| *Seelyosaurus guilelmiimperatori* | Dames, 1895 [39] |
| *Meyerasaurus victor* | Smith & Vincent, 2010 [2] |
| *Hauffiosaurus zanoni* | Vincent et al, 2011 [48] |
| *Atychodracon megacephalus* | Smith, 2015 [18] |
| *Thalassiodracon hawkinsii* | Smith, 2007 [34] |
| *Eoplesiosaurus antiquior* | Benson et al, 2012 [15] |
| *Hauffiosaurus tomistomimus* | Benson et al, 2011 [8] |
| *Microcleidus brachypterygius* | Huene, 1923 [49] |
| *Brancasaurus brancai* | Sachs et al, 2016 [17] |
| *Cryptoclidus eurymerus* | Andrews, 1910 [50] |
| *Nichollssaura borealis* | Druckenmiller & Russell, 2008 [42] |
| *Dolichorhynchops osborni* | Willison, 1907 [43] |
| *Wapuskanectes betsynichollsae* | Henderson, 2024 [45] |

### Proxy for Volume.xlsx

| | |
|---|---|
| *Abyssosaurus nataliae* | Berezin, 2011 [26]; Berezin, 2019 [27] |
| *Albertonectes vanderveldei* | Kubo et al, 2012 [47] |
| *Aristonectes quiriquinensis* | Otero et al, 2014 [51]; Otero et al, 2018 [52] |
| *Cryptoclidus eurymerus* | Andrews, 1910 [50]; Richards, 2011 [40] |
| *Dolichorhynchops osborni* | Bonner, 1964 [11] |
| *Hydrotherosaurus alexandrae* | Welles, 1943 [12] |
| *Kronosaurus MCZ 1285* | Romer & Lewis, 1959 [53]; McHenry, 2009 [20] |
| *Liopleurodon ferox* | Linder, 1913 [31] |
| *Macroplata tenuiceps* | Ketchum & Smith, 2010 [19] |
| *Martinectes bonneri* | Adams, 1997 [54] |
| *Mauriciosaurus fernandezi* | Frey et al, 2017 [44] |
| *Meyerasaurus victor* | Smith & Vincent, 2010 [2] |

| | |
|---|---|
| *Microcleidus tournemirensis* | Bardet et al, 1999 [29] |
| *Monquirasaurus boyacensis* | Noè & Gómez-Pérez, 2022 [32] |
| *Peloneustes philarchus* | Linder, 1913 [31] |
| *Pliosaurus funkei* | Knutsen et al, 2012 [55] |
| *Pliosaurus cf.kevani* | Tarlo, 1959 [56] |
| *Sachicasaurus vitae* | Páramo-Fonseca et al, 2018 [3] |
| *Seelyosaurus guilelmiimperatori* | Sachs et al, 2025 [57] |
| *Stenorhynchosaurus munozi* | Páramo-Fonseca et al, 2016 [36] |
| *Styxosaurus SDSM 451* | Welles & Bump, 1949 [9] |
| *Thalassomedon haningtoni* | Welles, 1943 [12] |
| *Vegasaurus molyi* | O'Gorman et al, 2015 [46] |

### raw data for size evolution_all.xlsx

| | |
|---|---|
| *Archaeonectrus rostratus* | Owen, 1865 [35] |
| *Attenborosaurus conybeari* | Benson et al, 2012 [15] |
| *Atychodracon megacephalus* | Smith, 2007 [34] |
| *Avalonnectes arturi* | Benson et al, 2012 [15] |
| *Brachauchenius lucasi* | McHenry, 2009 [20] |
| *Brancasaurus brancai* | Sachs et al, 2016 [17] |
| *Callawayasaurus colombiensis* | Welles, 1963 [25] |
| *Colymbosaurus megadeirus* | Benson & Bowdler, 2014 [58] |
| *Elasmosaurus platyurus* | Welles, 1952 [13] |
| *Eoplesiosaurus antiquior* | Benson et al, 2012 [15] |
| *Eretmosaurus rugosus* | Owen, 1865 [35] |
| *Fluvionectes sagecrensis* | Campbell et al, 2021 [59] |
| *Hauffiosaurus longirostris* | Benson et al, 2012 [15] |
| *Hauffiosaurus tomistomimus* | Benson et al., 2011 [8] |
| *Hauffiosaurus zanoni* | Vincent et al, 2010 [48] |
| *Kaiwhekea katiki* | Cruickshank & Fordyce, 2002 [4] |
| *Lindwurmia thiuda* | Vincent & Storrs, 2019 [60] |
| *Luskhan itilensis* | Fischer et al, 2023 [21] |
| *Microcleidus brachypterygius* | Benson et al, 2012 [15] |
| *Microcleidus homalospondylus* | Benson et al, 2012 [15] |
| *Morenosaurus stocki* | Welles, 1943 [12] |
| *Nichollssaura borealis* | Henderson, 2024 [45] |
| *Plesiosaurus dolichodeirus* | Storrs, 1997 [38] |
| *Rhomaleosaurus cramptoni* | Smith, 2007 [34] |
| *Rhomaleosaurus zetlandicus* | Benson et al, 2012 [15] |
| *Styxosaurus rezaci* | Welles, 1970 [61] |

| | |
|---|---|
| *Styxosaurus sp.* | Welles, 1952 [13] |
| *Tatenectes laramiensis* | O'Keefe et al, 2011 [41] |
| *Thalassiodracon hawkinsii* | Smith, 2007[34] |
| *Trinacromerum bentonianum* | Thurmond, 1968 [6] |
| *Wapuskanectes betsynichollsae* | Henderson, 2024 [45] |
| *Gronausaurus wegneri* | Hampe, 2013 [62] |
| *Libonectes morgani* | Gutarra et al, 2022 [63] |
| *Manemergus anguirostris* | Gutarra et al, 2022 [63] |
| *Plesiopterys wildi* | Gutarra et al, 2022 [63] |
| *Brachauchenius MNA V9433* | Albright III et al, 2007 [64] |
| *Colymbosaurus svalbardensis* | Roberts et al, 2017 [65] |
| *Dolichorhynchops herschelensis* | Sato, 2005 [66] |
| *Eopolycotylus rankini* | Albright III et al, 2010 [67] |
| *Futabasaurus suzukii* | Sato et al, 200 [68] |
| *Jucha squalea* | Fischer et al, 2020 [69] |
| *Leptocleidus superstes* | Kear & Barrett, 2011 [70] |
| *Makhaira rossica* | Fischer et al, 2015 [71] |
| *Muraenosaurus leedsii* | Andrews, 1910 [50] |
| *Nakonanectes bradti* | Serratos et al, 2017 [28] |
| *Ophthalmothule cryostea* | Roberts et al, 2020 [5] |
| *Palmulasaurus quadratus* | Albright III et al, 2010 [67] |
| *Pantosaurus striatus* | Wilhelm & O'Keefe, 2010 [72] |
| *Picrocleidus beloclis* | Andrews, 1910 [50] |
| *Pliosaurus andrewsi* | Andrews, 1913 [73] |
| *Rhomaleosaurus thorntoni* | Smith & Benson, 2015 [74] |
| *Scalamagnus tropicensis* | McKean, 2012 [75] |
| *Thililua longicollis* | Bardet et al, 2003 [7] |
| *Tricleidus seeleyi* | Andrews, 1910 [50] |
| *Westphaliasaurus simonsensii* | Schwermann & Sander, 2011 [76] |
| *Djupedalia engeri* | Knutsen et al, 2012 [77] |
| *Plesiosaurus mansellii* | Hulke, 1870 [78] |
| *Aphrosaurus furlongi* | O'Gorman, 2020 [79] |
| *Cardiocorax mukulu* | Aráujo et al, 2015 [80] |
| *Chubutinectes carmeloi* | O'Gorman et al, 2023 [81] |
| *Eurycleidus arcuatus* | Smith, 2007 [34] |
| *Kawanectes lafquenianum* | O'Gorman, 2019 [82] |
| *Plesiopharos moelensis* | Puèrtolas-Pascual et al, 2021 [83] |
| *Simolestes vorax* | Andrews, 1913 [73] |
| *Zarafasaura oceanis* | Lomax & Wahl, 2013 [84] |
| *Arminisaurus schuberti* | Sachs & Kear, 2017 [85] |
| *Eardasaurus powelli* | Ketchum & Benson, 2022 [86] |

| | |
|---|---|
| *Franconiasaurus brevispinus* | Sachs et al, 2024 [87] |
| *Morturneria seymourensis* | Lester, 2019 [88] |
| *Pahasapasaurus haasi* | Schumacher, 2007 [89] |
| *Pliosaurus brachyspondylus* | Tarlo, 1959 [56] |
| *Pliosaurus irgisensis* | Storrs et al, 2000 [90] |
| *Pliosaurus rossicus* | Halstead, 1971 [91] |
| *Marmornectes candrewi* | Ketchum & Benson, 2011 [92] |
| *Terminonatator ponteixensis* | Sato, 2003 [93] |
| *Umoonasaurus demoscyllus* | Kear et al, 2006 [94] |
| *Spitrasaurus spp* | Knutsen et al, 2012b [95] |
| *Aristonectes parvidens* | O'Gorman, 2015 [96] |
| *Edgarosaurus muddi* | Druckenmiller, 2002 [14] |
| *Pliosaurus almanzaensis* | O'Gorman et al, 2018 [97] |
| *Pliosaurus brachydeirus* | Knutsen, 2012 [98] |
| *Pliosaurus carpenteri* | Sassoon et al, 2012 [99] |
| *Pliosaurus kevani* | Benson et al, 2013 [100] |
| *Pliosaurus macromerus* | Owen, 1869 [101] |
| *Pliosaurus westburyensis* | Sassoon et al, 2012 [99] |
| *Serpentisuchops pfisterae* | Persons et al, 2022 [24] |
| *Styxosaurus snowii* | Sachs et al, 2018 [102] |
| *Thaumatodracon wiedenrothi* | Smith & Araújo, 2017 [30] |

[11] Orville Bonner. An osteological study of *Nyctosaurus* and *Trinacromerum* with a description of a new species of *Nyctosaurus*. Master's thesis, Fort Hays State University, 1964.

[12] Samuel Paul Welles. Elasmosaurid plesiosaurs with description of new material from california and colorado. *Memoirs of the University of California*, 13:125–254, 1943.

[13] Samuel Paul Welles. *A review of the North American Cretaceous elasmosaurs*. University of California Press, 1952.

[14] Pat S Druckenmiller. Osteology of a new plesiosaur from the lower cretaceous (albian) thermopolis shale of montana. *Journal of Vertebrate Paleontology*, 22(1):29–42, 2002.

[15] Roger B. J. Benson, Mark Evans, and Patrick S. Druckenmiller. High diversity, low disparity and small body size in plesiosaurs (reptilia, sauropterygia) from the triassic–jurassic boundary. *PLoS ONE*, 7(3):e31838, mar 2012.

[16] F Robin O'keefe. Preliminary description and phylogenetic position of a new plesiosaur (reptilia: Sauropterygia) from the toarcian of holzmaden, germany. *Journal of Paleontology*, 78(5):973–988, 2004.

[17] Sven Sachs, Jahn J. Hornung, and Benjamin P. Kear. Reappraisal of europe's most complete early cretaceous plesiosaurian: *Brancasaurus brancai* wegner, 1914 from the "wealden facies" of germany. *PeerJ*, 4:e2813, dec 2016.

[18] AS Smith. Reassessment of *'Plesiosaurus' megacephalus* (sauropterygia: Plesiosauria) from the triassic-jurassic boundary, UK. *Palaeontologia Electronica*, 2015.

[19] Hilary F. Ketchum and Adam S. Smith. The anatomy and taxonomy of *Macroplata tenuiceps* (sauropterygia, plesiosauria) from the hettangian (lower jurassic) of warwickshire, united kingdom. *Journal of Vertebrate Paleontology*, 30(4):1069–1081, jul 2010.

[20] Colin Richard McHenry. *Devourer of gods: the palaeoecology of the Cretaceous pliosaur Kronosaurus queenslandicus*. PhD thesis, University of Newcastle, 2009.

[21] Valentin Fischer, Roger BJ Benson, Nikolay G Zverkov, Maxim S Arkhangelsky, Ilya M Stenshin, Gleb N Uspensky, and Natalya E Prilepskaya. Anatomy and relationships of the bizarre early cretaceous pliosaurid luskhan itilensis. *Zoological Journal of the Linnean Society*, 198(1):220–256, 2023.

[22] F Robin O'Keefe. The evolution of plesiosaur and pliosaur morphotypes in the plesiosauria (reptilia: Sauropterygia). *Paleobiology*, 28(1):101–112, 2002.

[23] David Seymour Brown. The english upper jurassic plesiosauroidea (reptilia) and a review of the phylogeny and classification of the plesiosauria. 1981.

[31] Hermann Linder. Beiträge zur kenntnis der plesiosaurier-gattungen *Peloneustes* und *Pliosaurus*. *Geologische und Palaeontologishe Abhandlungen*, 1913.

[32] Leslie F. Noè and Marcela Gómez-Pérez. Giant pliosaurids (sauropterygia; plesiosauria) from the lower cretaceous peri-gondwanan seas of colombia and australia. *Cretaceous Research*, 132:105122, apr 2022.

[33] Ramon Nagesan. Neck mobility of the plesiosaur *Nichollsaura borealis*. Master's thesis, University of Calgary, 2017.

[34] Adam Stuart Smith. *Anatomy and systematics of the Rhomaleosauridae (Sauropterygia: Plesiosauria)*. PhD thesis, 2007.

[35] Owen. Monograph of the fossil reptilia of the liassic formations. part first. sauropterygia. *Monographs of the Palaeontographical Society*, 17(75):1–40, 1865.

[36] María Eurídice Páramo-Fonseca, Marcela Gómez-Pérez, Leslie F. Noè, and Fernando Etayo-Serna. *Stenorhynchosaurus munozi*, gen. et sp. nov. a new pliosaurid from the upper barremian (lower cretaceous) of villa de leiva, colombia, south america. *Revista de la Academia Colombiana de Ciencias Exactas, Físicas y Naturales*, 40(154):84, March 2016.

[37] Peggy Vincent, Didier Poncet, Anthony Rard, Jean-Pierre Robin, and Rémi Allemand. New remains of *Liopleurodon* (reptilia, plesiosauria) from the middle jurassic of western france and paedomorphosis within pliosaurids. *Palaeontologia Electronica*, 2024.

[38] Glenn W Storrs. Morphological and taxonomic clarification of the genus plesiosaurus. In *Ancient marine reptiles*, pages 145–190. Elsevier, 1997.

[39] Wilhelm Dames. Die plesiosaurier der süddeutschen liasformation. 1895.

[40] Courtney D Richards. Plesiosaur body shape and its impact on hydrodynamic properties, 2011.

[41] F. Robin O'Keefe, Hallie P. Street, Benjamin C. Wilhelm, Courtney D. Richards, and Helen Zhu. A new skeleton of the cryptoclidid plesiosaur *Tatenectes laramiensis* reveals a novel body shape among plesiosaurs. *Journal of Vertebrate Paleontology*, 31(2):330–339, mar 2011.

[42] Patrick S. Druckenmiller and Anthony P. Russell. Skeletal anatomy of an exceptionally complete specimen of a new genus of plesiosaur from the early cretaceous (early albian) of northeastern alberta, canada. *Palaeontographica Abteilung A*, 283(1–3):1–33, March 2008.

[43] Samuel Wendell Williston. *North American plesiosaurs: Part I*, volume 2. Field Columbian Museum, 1907.

[50] Charles William Andrews. *A descriptive catalogue of the marine reptiles of the Oxford clay: Based on the Leeds Collection in the British Museum (Natural History), London*, volume 1. order of the Trustees, 1910.

[57] Sven Sachs, Daniel Madzia, Miguel Marx, Aubrey J. Roberts, Oliver Hampe, and Benjamin P. Kear. The osteology, taxonomy, and phylogenetic placement of *Seeleyosaurus guilelmiimperatoris* (plesiosauroidea, microcleididae) from the lower jurassic posidonia shale of germany. *The Anatomical Record*, February 2025.

[58] Roger B. J. Benson and Timothy Bowdler. Anatomy of *Colymbosaurus megadeirus* (reptilia, plesiosauria) from the kimmeridge clay formation of the u.k., and high diversity among late jurassic plesiosauroids. *Journal of Vertebrate Paleontology*, 34(5):1053–1071, July 2014.

[59] James A. Campbell, Mark T. Mitchell, Michael J. Ryan, and Jason S. Anderson. A new elasmosaurid (sauropterygia: Plesiosauria) from the non-marine to paralic dinosaur park formation of southern alberta, canada. *PeerJ*, 9:e10720, feb 2021.

[60] Peggy Vincent and Glenn W. Storrs. *Lindwurmia*, a new genus of plesiosauria (reptilia: Sauropterygia) from the earliest jurassic of halberstadt, northwest germany. *The Science of Nature*, 106(1-2), jan 2019.

[61] SP Welles. The longest neck in the ocean. *University of Nebraska News, Museum Notes*, 1970.

[73] Charles William Andrews. *A descriptive catalogue of the marine reptiles of the Oxford clay: Based on the Leeds Collection in the British Museum (Natural History), London*, volume 2. order of the Trustees, 1913.

[74] Adam S. Smith and Roger B.J. Benson. Osteology of *Rhomaleosaurus thorntoni* (sauropterygia: Rhomaleosauridae) from the lower jurassic (toarcian) of northamptonshire, england. *Monographs of the Palaeontographical Society*, 168(642):1–40, oct 2014.

[75] Rebecca Schmeisser McKean. A new species of polycotylid plesiosaur (reptilia: Sauropterygia) from the lower turonian of utah: Extending the stratigraphic range of dolichorhynchops. *Cretaceous Research*, 34:184–199, April 2012.

[76] Leonie Schwermann. Osteologie und phylogenie von *Westphaliasaurus simonsensii*: Ein neuer plesiosauride (sauropterygia) aus dem unteren jura (pliensbachium) von sommersell (kreis höxter), nordrhein-westfalen, deutschland. *Geol. Paläont. Westf.*, 79:1, 2011.

[77] Espen M Knutsen, Patrick S Druckenmiller, and Jørn H Hurum. A new plesiosauroid (reptilia: Sauropterygia) from the agardhfjellet formation (middle volgian) of central spitsbergen, norway. *Norwegian Journal of Geology*, 92:213–234, 2012.

[78] JW Hulke. Note on some plesiosaurian remains obtained by jc mansel, esq., fgs, in kimmerldge bay, dorset. *Quarterly Journal of the Geological Society*, 26(1-2):611–622, 1870.

[79] Jose P O'Gorman. Elasmosaurid phylogeny and paleobiogeography, with a reappraisal of *Aphrosaurus furlongi* from the maastrichtian of the moreno formation. *Journal of Vertebrate Paleontology*, 39(5):e1692025, 2019.

[80] R. Araújo, M.J. Polcyn, J. Lindgren, L.L. Jacobs, A.S. Schulp, O. Mateus, A. Olímpio Gonçalves, and M.-L. Morais. New aristonectine elasmosaurid plesiosaur specimens from the early maastrichtian of angola and comments on paedomorphism in plesiosaurs. *Netherlands Journal of Geosciences - Geologie en Mijnbouw*, 94(1):93–108, feb 2015.

[81] José P. O'Gorman, Ana Paula Carignano, Lydia Calvo-Marcilese, and Juan Pablo Pérez Panera. A new elasmosaurid (sauropterygia, plesiosauria) from the upper levels of the la colonia formation (upper maastrichtian), chubut province, argentina. *Cretaceous Research*, 152:105674, December 2023.

[82] JosÉ Patricio O'Gorman. First record of *Kawanectes lafquenianum* (plesiosauria, elasmosauridae) from the la colonia formation of argentina, with comments on the mandibular morphology of elasmosaurids. *Alcheringa: An Australasian Journal of Palaeontology*, 44(1):176–193, nov 2019.

[83] Eduardo Puértolas-Pascual, Miguel Marx, Octávio Mateus, André Saleiro, Alexandra Fernandes, João Marinheiro, Carla Tomás, and Simão Mateus. A new plesiosaur from the lower jurassic of portugal and the early radiation of plesiosauroidea. *Acta Palaeontologica Polonica*, 66, 2021.

[84] Dean Lomax and William Wahl. A new specimen of the elasmosaurid plesiosaur *Zarafasaura oceanis* from the upper cretaceous (maastrichtian) of morocco. *Paludicola*, 9:97–109, 10 2013.

[85] Sven Sachs and Benjamin P. Kear. A rare new pliensbachian plesiosaurian from the amaltheenton formation of bielefeld in northwestern germany. *Alcheringa: An Australasian Journal of Palaeontology*, 42(4):487–500, sep 2017.

[86] Hilary Ketchum and Roger Benson. A new pliosaurid from the oxford clay formation of oxfordshire, UK. *Acta Palaeontologica Polonica*, 67, 2022.

[87] Sven Sachs, Stefan Eggmaier, and Daniel Madzia. Exquisite skeletons of a new transitional plesiosaur fill gap in the evolutionary history of plesiosauroids. *Frontiers in Earth Science*, 12, February 2024.

[88] Elizabeth Lester. Description and histology of a small-bodied elasmosaur and discription of mortuneria seyemourensis postcranium. 2019.

[89] Bruce A Schumacher. A new polycotylid plesiosaur (reptilia; sauropterygia) from the greenhorn limestone (upper cretaceous; lower upper cenomanian), black hills, south dakota. *The Geological Society of America Special Paper*, 2007.

[101] Richard Owen. *Monographs on the British Fossil Reptilia from the Kimmeridge Clay. No. III, Containing Pliosaurus Grandis, Pl. Trochanterius, and Pl. Portlandicus. Pages 1–12; Plates I—IV*, volume 22. Taylor & Francis, 1869.

[102] Sven Sachs, Johan Lindgren, and Benjamin P. Kear. Reassessment of the *Styxosaurus snowii* (williston, 1890) holotype specimen and its implications for elasmosaurid plesiosaurian interrelationships. *Alcheringa: An Australasian Journal of Palaeontology*, 42(4):560–574, October 2018.
